# Monoallelically-expressed Noncoding RNAs form nucleolar territories on NOR-containing chromosomes and regulate rRNA expression

**DOI:** 10.1101/2022.07.04.498693

**Authors:** Qinyu Hao, Minxue Liu, Swapna Vidhur Daulatabad, Saba Gaffari, Rajneesh Srivastava, You Jin Song, Shivang Bhaskar, Anurupa Moitra, Hazel Mangan, Elizabeth Tseng, Rachel B. Gilmore, Susan M. Freier, Xin Chen, Chengliang Wang, Sui Huang, Stormy Chamberlain, Hong Jin, Jonas Korlach, Brian McStay, Saurabh Sinha, Sarath Chandra Janga, Supriya G. Prasanth, Kannanganattu V. Prasanth

## Abstract

Out of the several hundred copies of *rRNA* genes that are arranged in the nucleolar organizing regions (NOR) of the five human acrocentric chromosomes, ∼50% remain transcriptionally inactive. NOR-associated sequences and epigenetic modifications contribute to differential expression of rRNAs. However, the mechanism(s), controlling the dosage of active versus inactive *rRNA* genes in mammals is yet to be determined. We have discovered a family of ncRNAs, SNULs (<u>S</u>ingle <u>NU</u>cleolus <u>L</u>ocalized RNA), which form constrained sub-nucleolar territories on individual NORs and influences rRNA expression. Individual members of the SNULs monoallelically associate with specific NOR-containing chromosome. SNULs share sequence similarity to pre-rRNA and localize in the sub-nucleolar compartment with pre-rRNA. Finally, SNULs control rRNA expression by influencing pre-rRNA sorting to the DFC compartment and pre-rRNA processing. Our study discovered a novel class of ncRNAs that by forming constrained nucleolar territories on individual NORs contribute to rRNA expression.

## INTRODUCTION

The nucleolus is the most well-characterized non-membranous nuclear domain, where ribosome biogenesis and maturation takes place and is formed around the nucleolus organizer regions (NORs) ^1^. NORs are comprised of rRNA gene tandem arrays, and in human cells, they are located on the short arms (p-arm) of the five acrocentric chromosomes (Chrs. 13,14,15, 21 & 22) ^2^. Human cells contain >400 copies of rRNA (18S/28S/5.8S) genes, yet only ∼50% of the copies are transcriptionally active ^3^. The expression of rRNA genes is tightly controlled during physiological processes, such as cellular development by epigenetic mechanisms ^2, 4–6^. However, the mechanism that precisely maintains the dosage of active versus inactive rRNA genes within a cell is yet to be determined.

The nucleolus harbors a diverse set of small and long noncoding RNAs (ncRNAs), which play crucial roles in organizing the nucleolar genome as well as regulating rRNA expression ^7, 8^. For example, the intergenic spacer (IGS) between rRNA genes encodes several ncRNAs, such as pRNA, PAPAS, and PNCTR, which modulate rRNA expression and nucleolus organization ^7–10^. Recent studies have reported that ncRNAs, including SLERT, LoNa, AluRNAs and the LETN lncRNAs modulate nucleolus structure and rRNA expression via independent mechanisms ^11–14^. Collectively, these studies underscore the importance of ncRNAs in controlling key nucleolus functions thus contributing to cellular homeostasis.

Besides the rDNA array, the remaining DNA sequences within the short arms of all five NOR-containing acrocentric chromosomes are highly repetitive in nature and share higher levels of sequence similarities ^2, 4, 5, 15–17^. As a result, insights into novel genes and/or regulatory elements located within the p-arms are limited. Analyses of small regions located adjacent to NORs revealed that they code for lncRNAs ^15, 18^, indicating that the p-arms of the NOR-containing chromosomes harbor ncRNA genes, those could modulate key nucleolar functions.

In the present study, we have identified a novel family of ncRNAs: SNULs, which likely originate from the p-arms of acrocentric chromosomes and form allele-specific constrained sub-nucleolar territories on the NOR-containing chromosomes. SNUL-1 RNA displays high sequence similarity to pre-rRNA. Significantly, our studies revealed that the SNUL family of ncRNAs contribute to rRNA expression. Thus, our study unraveled the existence of a novel family of ncRNAs that display monoallelic coating/association on the autosomal segments of NOR-containing p-arms for modulating rRNA expression.

## RESULTS

### SNUL-1 RNA forms a distinct territory within the nucleolus

In a screen to identify RNAs with distinct cellular distribution ^19^, we identified a unique probe with ∼600 nucleotides (Table S1), which preferentially hybridized to an RNA species that formed a cloud/territory within the nucleolus in a broad spectrum of human cell lines (Figures 1a-b & S1a-b). Unlike other nucleolus-resident RNAs, which is homogenously distributed in all the nucleoli within a cell, this RNA cloud decorated one nucleolus of several nucleoli per nucleus in most of the diploid or near-diploid cells (Figure 1a-c & S1a; embryonic stem cell [WA09], fibroblasts [WI-38, IMR-90, and MCH065], epithelial cell [hTERT-RPE-1], and lymphocyte [GM12878]). We therefore named the RNA as <u>S</u>ingle <u>NU</u>cleolus <u>L</u>ocalized RNA-1 (SNUL-1). Strikingly, cancer cell lines displayed varied numbers of the SNUL-1 territories per nucleus (ranging from 1 to 4 SNUL-1 clouds/cell), though the number of SNUL-1 cloud/cell remained fixed for a particular cell line (Figure 1c). The SNUL-1 cloud was well-preserved even in biochemically isolated nucleoli (Figure 1d), indicating that SNUL-1 associates with integral components of the nucleolus.

**Fig. 1.**
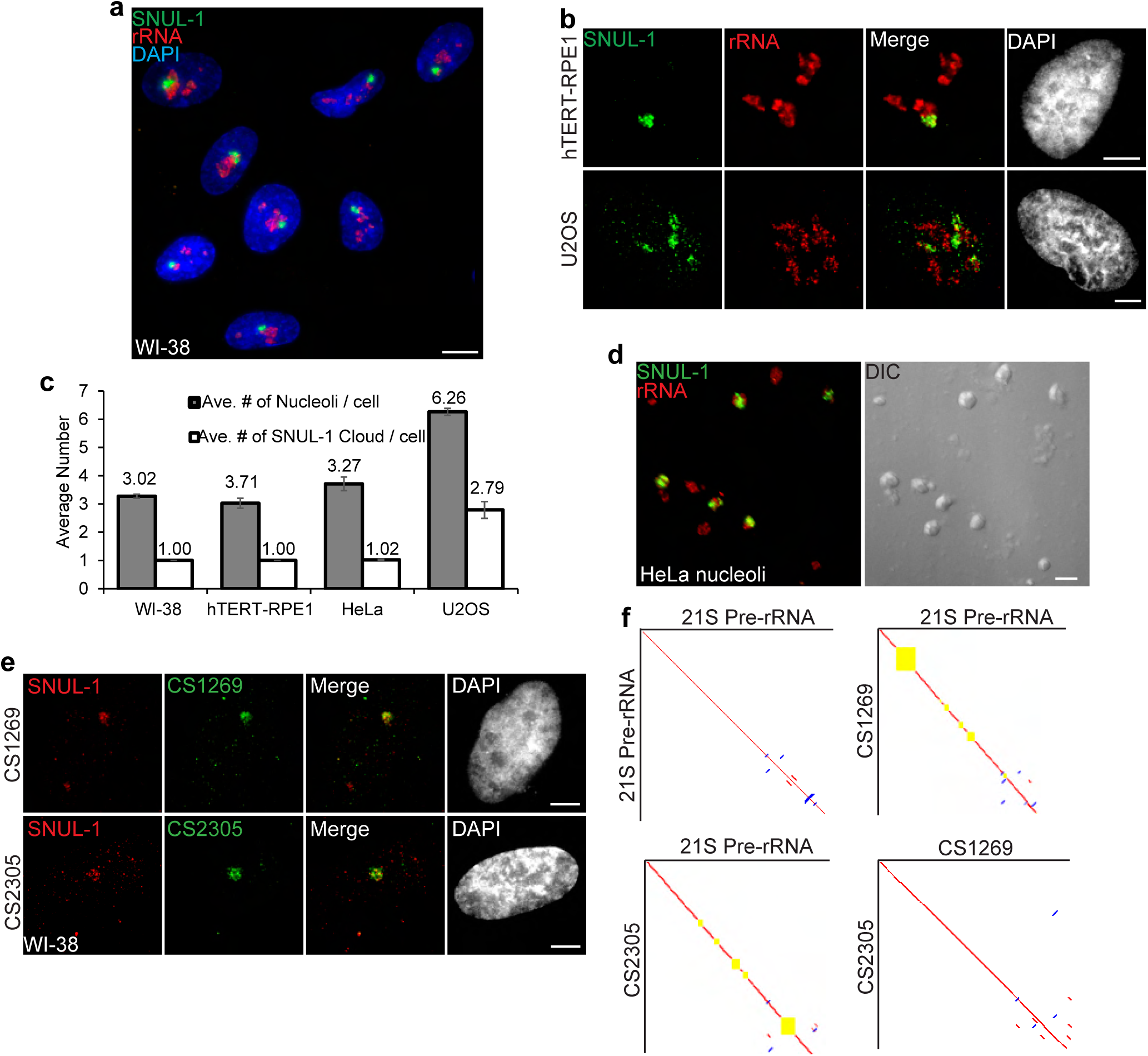
SNUL-1 forms RNA clouds in human cell lines. **a**, RNA-FISH of SNUL-1 (green) in WI-38 cells. Nucleoli are visualized by rRNA (red). **b**, RNA-FISH of SNUL-1 (green) in hTERT-RPE1 and U2OS cell lines. Nucleoli are visualized by rRNA (red). **c**, Graph depicting the average number of nucleoli/cell and the SNUL-1 clouds/cell in various cell lines. **d**, RNA-FISH of SNUL-1 (green) in biochemically isolated HeLa nucleoli marked by rRNA (red). Note that the distribution of SNUL-1 is preserved in the isolated nucleoli. **e**, RNA-FISH performed using probes designed from the SNUL-1 CSs (green) and SNUL-1 (red) Probe 1 in WI-38 cells. **f**, Pairwise sequence comparisons between 21S (pre-rRNA) and 21S, 21S and CS1269 (SNUL-1 CS), 21S and CS2305, and CS1269 and CS2305. Red lines indicate forward aligned regions, blue lines indicate reverse aligned regions, and yellow boxes indicate unaligned regions. All scale bars, 5µm.

### SNUL-1 constitutes a group of RNAs with sequence features resembling 21S pre-rRNA

The original double-stranded DNA probe (Probe 1; Figure S1c) that detected the SNUL-1 RNA cloud(s) was mapped to hg38-Chr17: 39549507-39550130 genomic region, encoding a lncRNA. However, other unique probes (non-overlapping with the probe-1 region) generated from the Chr17-encoded lncRNA failed to detect SNUL-1 RNA cloud (data not shown). Furthermore, BLAST-based analyses failed to align the Probe 1 sequence to any other genomic loci. Since a large proportion of the p-arms of nucleolus-associated NOR-containing acrocentric chromosomes is not yet annotated, we speculated that SNUL-1 could be transcribed from an unannotated genomic region from the acrocentric p-arms. RNA-FISH-based analyses revealed that a [CT]_20_ repeat and a 60- nucleotide overhang sequence within the original probe-1 was crucial for detecting the SNUL-1 cloud (Figures S1c-d; probe 4), implying that the [AG] repeats along with unique sequence beyond the repeat contributes to the hybridization specificity and localization of SNUL-1.

During the screen, we identified another single-stranded oligonucleotide probe that shared ∼73% sequence similarity to the SNUL-1 probe 4 detected an additional RNA cloud in the nucleolus, but was distinct from the SNUL-1 cloud (Figure S1e-f). We named this RNA as SNUL-2. The probe that hybridized to SNUL-2 also contained an imperfect [CT]-rich region (Figure S1f), suggesting that both SNUL-1 and SNUL-2 RNAs contain [AG] repeats. Based on this, we propose that SNUL-1 and SNUL-2 are members of a novel RNA family, which form non-overlapping constrained territories within the nucleolus.

In order to identify the full-length SNUL-1 sequence, we isolated rRNA-depleted total RNA from biochemically purified nucleoli (Figure 1d) and performed targeted long-read Iso-Sequencing (Iso-Seq; Single Molecule Real-Time Sequencing by PACBIO) using the SNUL-1 Probe 4 as a bait for the initial RNA pull down (Figure S1g; please see methods for details). Top ranked full-length isoforms with high binding affinity with SNUL-1 Probe 4 were picked as potential SNUL-1 candidate sequences (CSs). We identified potential SNUL-1 candidates that were supported by both Iso-Seq and an independent unbiased Nanopore long-read sequencing of nucleolus-enriched RNAs (Figure S1g). Comparison between the Iso-Seq SNUL-1 CSs and the independently built consensus sequences by Nanopore reads revealed a ∼100% identity, thus confirming the presence of SNUL-1 full-length candidates in the nucleolus. Furthermore, individual members of the SNUL-1 CS RNAs were localized within the SNUL-1 RNA territory (Figures 1e & S1h). Members of the SNUL-1 CS RNAs, though enriched with the same SNUL-1 cloud, did not display complete co-localization (Figures S1i-j). The full-length SNUL-1 CS RNAs identified by both iso-seq and nanopore seq. analyzes contained defined 5’ and 3’ends, ranges in length from 1.9kb to 3.1kb and displayed high levels of sequence similarity between each other (>90%) (Figures 1f & S1k). Furthermore, comprehensive statistical analyses support the inference that SNUL-1 CSs constituted a group of independent RNAs, sharing high levels of sequence similarity (Figure S1l & Tables S2 & S3, see Materials and Methods for details of the analyses). Interestingly, the individual members of the SNUL-1 CSs showed ∼80% sequence similarity to 21S pre-ribosomal RNA (pre- rRNA), which is an intermediate pre-rRNA, consisting of 18S rRNA and partial internal transcribed spacer 1 (ITS1) (Figures 1f, S1k & S1m).

In the nucleolus, both SNUL-1 and pre-rRNA (detected by the probe hybridizing to the internal transcribed region [ITS1] of pre-rRNA) distributed mostly non-overlapping regions, as observed by super resolution-structured illumination microscopy (SR-SIM) imaging (Figure S1n). In addition, depletion of SNULs using modified DNA antisense oligonucleotides did not reduce the levels of pre-rRNA (Figure S1o). Based on these results, we conclude that SNUL-1 represents a group of RNA species showing sequence similarities to 21S pre-rRNA and form a constrained single territory within the nucleolus.

### RNA polymerase I-transcribed SNUL-1 is enriched at the DFC sub-nucleolar region

In the nucleolus, RNA Pol I transcription machinery is clustered in the fibrillar center (FC; marked by UBF or RNA polymerase 1 [RPA194]), allowing the transcription to happen at the outer boundary of FC ^20^. Nascent pre-rRNAs are co-transcriptionally sorted into the dense fibrillar center (DFC; marked by fibrillarin [FBL]) located around FC for the early stages of pre-rRNA processing. The final steps of pre-rRNA processing and ribosome assembly take place in the granular component (GC; marked by B23) (Figure S2a). SR-SIM imaging revealed that SNUL-1 distributed across all the three sub- nucleolar compartments (Figures 2a & S2b) but preferentially enriched in the DFC region (higher Pearson’s correlation coefficient [PCC] between SNUL-1 and FBL [DFC marker] over RPA194 [FC marker]) (Figure 2a-b).

**Fig. 2.**
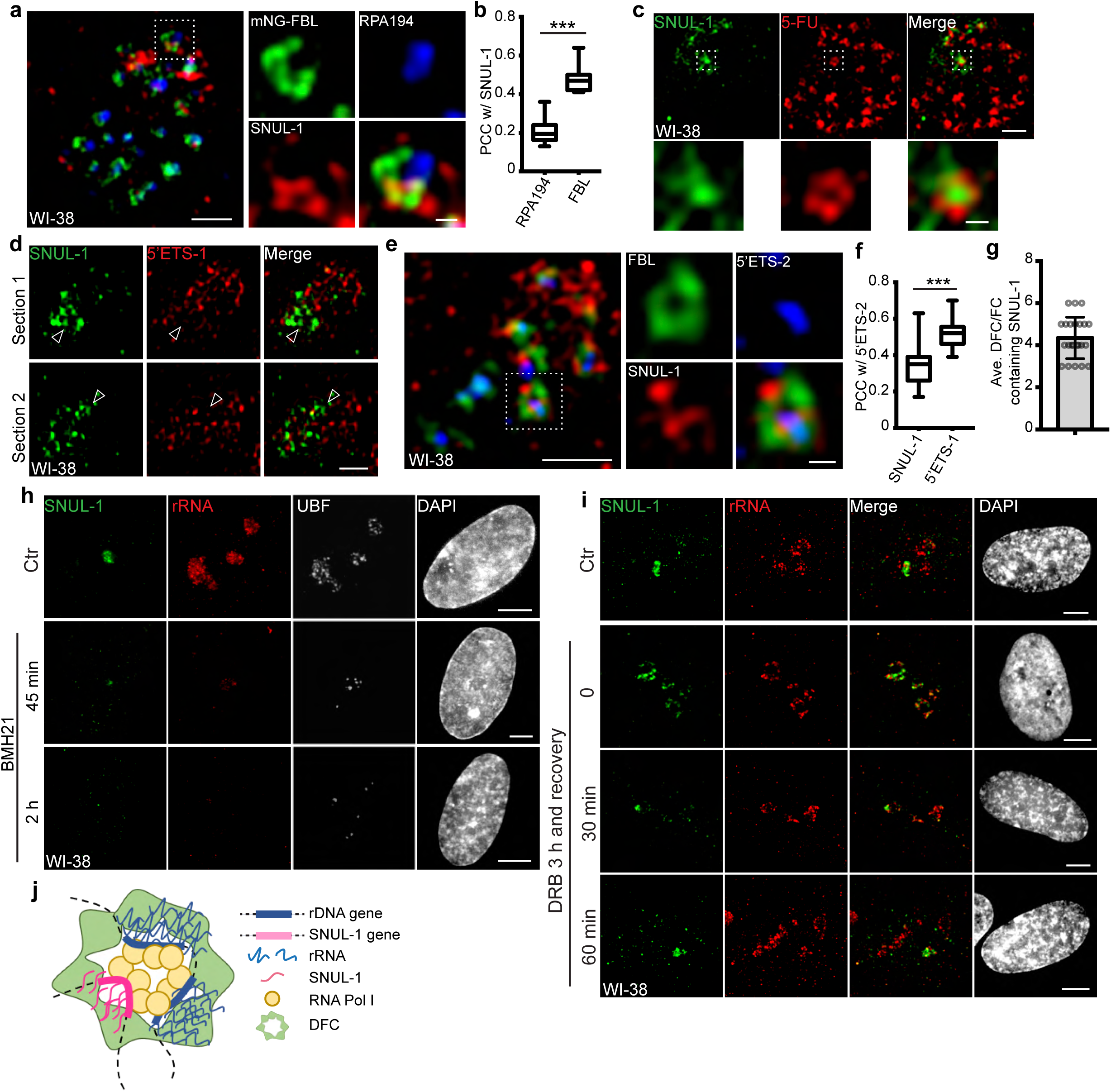
SNUL-1 is an RNA Pol I transcript and forms constrained nucleolar territory. **a**, Representative SIM image of the SNUL-1 (red) distribution relative to DFC/FC units in WI-38 cells. FC is marked by RPA194 (blue) and DFC is marked by mNeonGreen (NG)-FBL (green). Scale bars, 1 µm (main images) and 200nm (insets). **b**, Box plots showing Pearson’s correlation coefficients (PCCs) between SNUL-1 and either RPA194 (FC) or FBL (DFC). n = 16 and 11, respectively. Statistical analysis was performed using Mann-Whitney test. *p < 0.05, **p < 0.01, ***p < 0.001. Center line, median; box limits, upper and lower quartiles; whiskers, maximum or minimum of the data. **c**, Representative SIM image of the SNUL-1 (green) distribution relative to the 5-FU signal (red) in WI-38 cells. Nascent RNAs are metabolically labeled by 5 min of 5-FU pulse. Scale bars, 1 µm (main images) and 200nm (insets). **d**, two sections of SIM images from a single nucleolus showing relative distribution of SNUL-1 (green) and nascent pre-rRNA (marked by 5’ETS-1 probe) signals (red) in WI-38 cells. **e**, Representative SIM image of the SNUL-1 (red) distribution relative to DFC/FC unit and pre-rRNAs in WI-38 cells. DFC is marked by FBL (green) and pre-rRNAs (blue) are detected by 5’ETS-2 probe. Scale bars, 1 µm (main images) and 200nm (insets). **f**, Box plots showing the Pearson’s correlation coefficients (PCCs) between 5’ETS-2 signal and either SNUL-1 or 5’ETS-1 signal. n = 23 and 16, respectively. Statistical analysis was performed using Mann-Whitney test. *p < 0.05, **p < 0.01, ***p < 0.001. **g**, Graph depicting the average number of SNUL-1 positive DFC/FC units/nucleolus in WI-38 cells. Center line, median; box limits, upper and lower quartiles; whiskers, maximum or minimum of the data. **h**, Co-RNA-FISH and IF to detect SNUL-1 (green), rRNA (red) and UBF (white) in control and BMH21-treated WI-38 cells. Scale bars, 5 µm. **i**, RNA-FISH to detect SNUL-1 (green), rRNA (red) in control and DRB-treated WI-38 cells. For recovery after DRB treatment, the drug is washed off after 3 h of treatment and RNA-FISH is performed at 0, 30min and 60min timepoints during recovery. Scale bars, 5µm. **j**, Model showing the association of both SNUL-1 and rRNA in the same DFC/FC unit. DNA is counterstained with DAPI.

The SNUL-1 cloud associated with the transcriptionally active DFC/FC units as observed by the presence of 5-FU (fluro-uridine)-incorporated nascent RNA in SNUL-1- associated domains (Figure 2c). SNUL-1 positive regions within the nucleolus never completely overlapped with, but instead were located adjacent to pre-rRNA as well as rDNA signals, as observed by SR-SIM imaging (Figures 2d & S2c). However, both SNUL-1 and pre-rRNA co-existed but were not colocalized within an individual FC/DFC unit (Figures 2e-f & S2d). An individual nucleolus contains several dozens of FC/DFC units, with each unit containing 2-3 transcriptionally active rRNA genes ^21^. SNUL-1 co- occupied in ∼4 adjacent FC/DFC units within a single nucleolus along with pre-rRNA (Figure 2g) (n =22). The localization of SNUL-1 in multiple FC/DFC units along with the observed sequence variations between SNUL-1 CSs imply that SNUL-1 RNAs are transcribed by a family of genes located in 3-4 adjacent FC/DFC units and form a constrained sub-nucleolar territory.

SNUL-1 is transcribed by RNA Pol I, as cells treated with RNA Pol I inhibitors (BMH21 or low dose of Actinomycin D [ActD, 10 ng/ml]) showed reduced SNUL-1 levels (Figures 2h and S2e). In RNA Pol II-inhibited cells (flavopiridol and 5,6-Dichloro- 1-β-d-ribofuranosylbenzimidazole [DRB]), SNUL-1 re-localized to the nucleolar periphery along with rRNA (Figures 2i & S2f), and this alteration in the RNA distribution was reversible upon transcription reactivation (Figure 2i) ^22^. Together, these results indicated that SNUL-1 RNAs are transcribed by RNA Pol I in the FC/DFC region along with pre-rRNAs (Figure 2j).

### SNUL-1 RNA cloud associates with an NOR-containing chromosome

During mitosis, the SNUL-1 cloud was not observed from pro-metaphase to anaphase (Figure S3a). During late telophase/early G1, the nucleolus is formed around the active NORs ^23^. In the telophase/early G1 nuclei of multiple cell types, we observed a prominent single SNUL-1 cloud that co-localized with one of the several rRNA containing active NORs (see arrow in Figure S3a-b;). Late telophase or early G1 cells also showed a weak but distinct second SNUL-1 signal associating with another active NOR (please see arrowhead in Figure S3a-b). This result implies that SNUL-1 could be biallelically expressed, though at different levels during telophase/early G1 nuclei.

In the interphase nuclei, only one SNUL-1 cloud was observed that specifically associated with a single NOR-containing chromosome allele (Figure 3a). In this assay, the NOR-containing acrocentric chromosome arms were labeled by a probe hybridizing to the distal junction (DJ) regions, which are uniquely present on the p-arm of all the NOR-containing chromosomes^15, 18^. Further experiments revealed that in WI-38 interphase nuclei, including that of G1 cells, the SNUL-1 cloud specifically associated with one allele of Chr. 15. This was demonstrated by co-RNA and DNA-FISH, which detected SNUL-1 cloud and Chr. 15 markers, including Chr. 15 q-arm paint (Figures 3b-c & S3i), Chr. 15-specific centromere (15CEN; α-Satellite or 15p11.1-q11.1), and peri- centromeric Satellite III repeats (15Sat III repeats or 15p11.2) (Figure 4a). Monoallelic association of SNUL-1 to Chr. 15 was also confirmed in other cell lines, including in human primary fibroblasts (IMR-90 [lung], MCH065 [dermal]) and hTERT-immortalized near-diploid retinal pigment epithelial cells (hTERT-RPE1) (Figures S3c-d & 4c). In addition, we also observed that the SNUL-2 RNA cloud associated with one allele of NOR-containing Chr.13 (Figure S3e-f). Based on these results, we conclude that a unique subset of SNUL-like genes are present in each of the NOR-containing acrocentric chromosome arms, where each members of the SNUL RNA form a spatially constrained RNA territory on the p-arm of the particular chromosome allele.

**Fig. 3.**
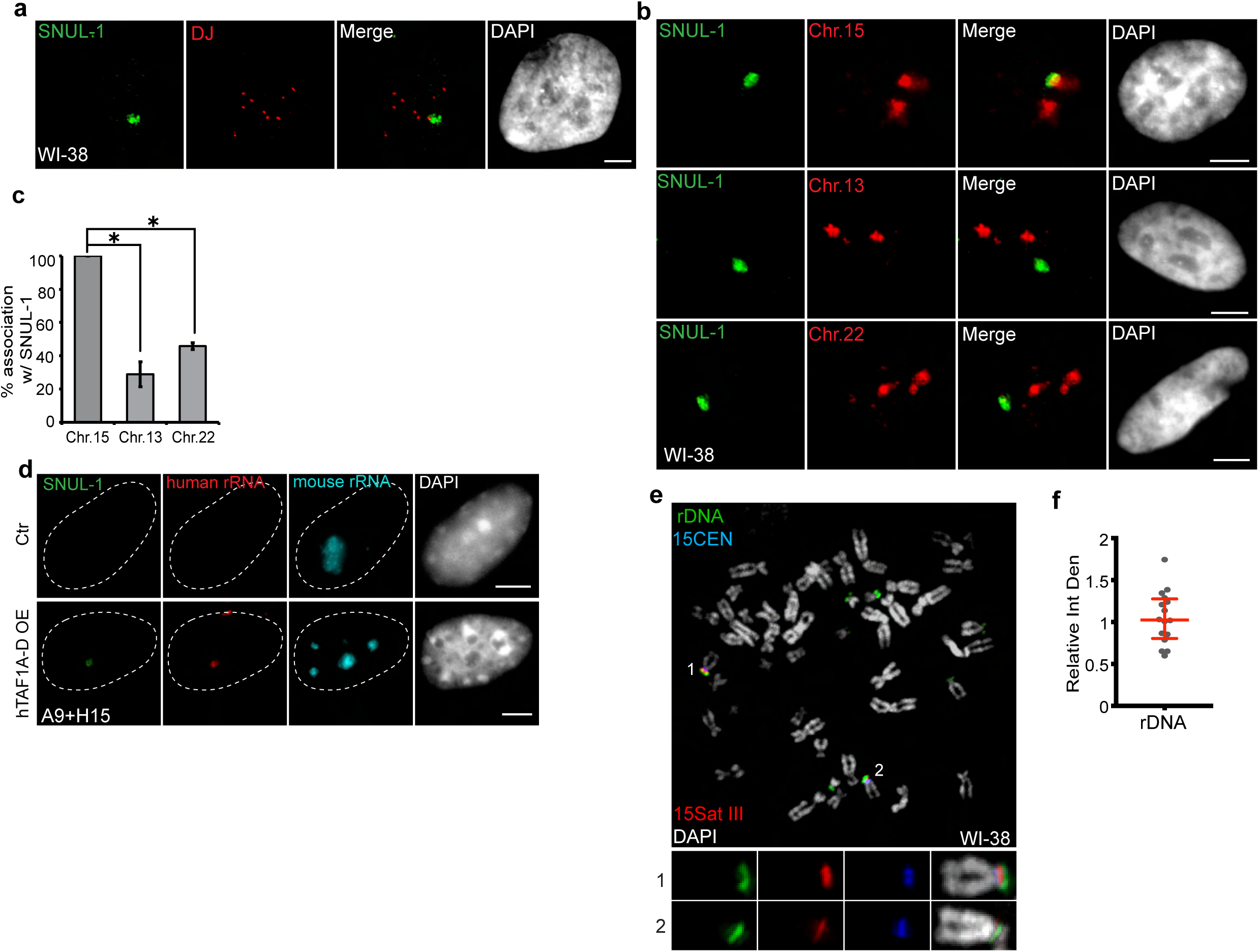
SNUL-1 is associated with the NOR of one Chr. 15 allele. **a**, DNA-RNA-FISH of SNUL-1 RNA (green) and distal junction (DJ) DNA (red) in WI-38 cells. **b**, DNA-RNA-FISH of SNUL-1 RNA and Chr. 15, Chr. 13, and Chr. 22 marked by probes painting the q-arms of the chromosomes in WI-38 cells. **c**, Quantification of the associa- tion rates between SNUL-1 and NOR containing chromosomes. Data are presented as Mean ± SD from biological triplicates. > 50 cells were counted for each of the biological repeats. Student’s unpaired two-tailed t-tests were performed. *p < 0.05. **d**, RNA-FISH to detect SNUL-1 (green), human rRNA (red) and mouse rRNA (blue) in control and hTAF1A-D overexpressed A9+H15 cells. Dotted lines mark the boundary of the nuclei. **e**, DNA-FISH showing rDNA and CEN15 and 15 Sat III contents on Chr. 15 in WI-38 metaphase chromosomes. The two alleles of Chr.15 are marked by 15Sat III and 15CEN, rDNA arrays are detected by a probe within the IGS region (See Fig. S1m). **f**, Relative integrated density of the two Chr. 15 rDNA arrays is calculated by dividing the measure- ment of the rDNA signal on the Chr. 15 with larger 15Sat III by that of the one on the Chr.15 with smaller 15Sat III. All scale bars, 5µm. DNA is counterstained with DAPI.

**Fig. 4.**
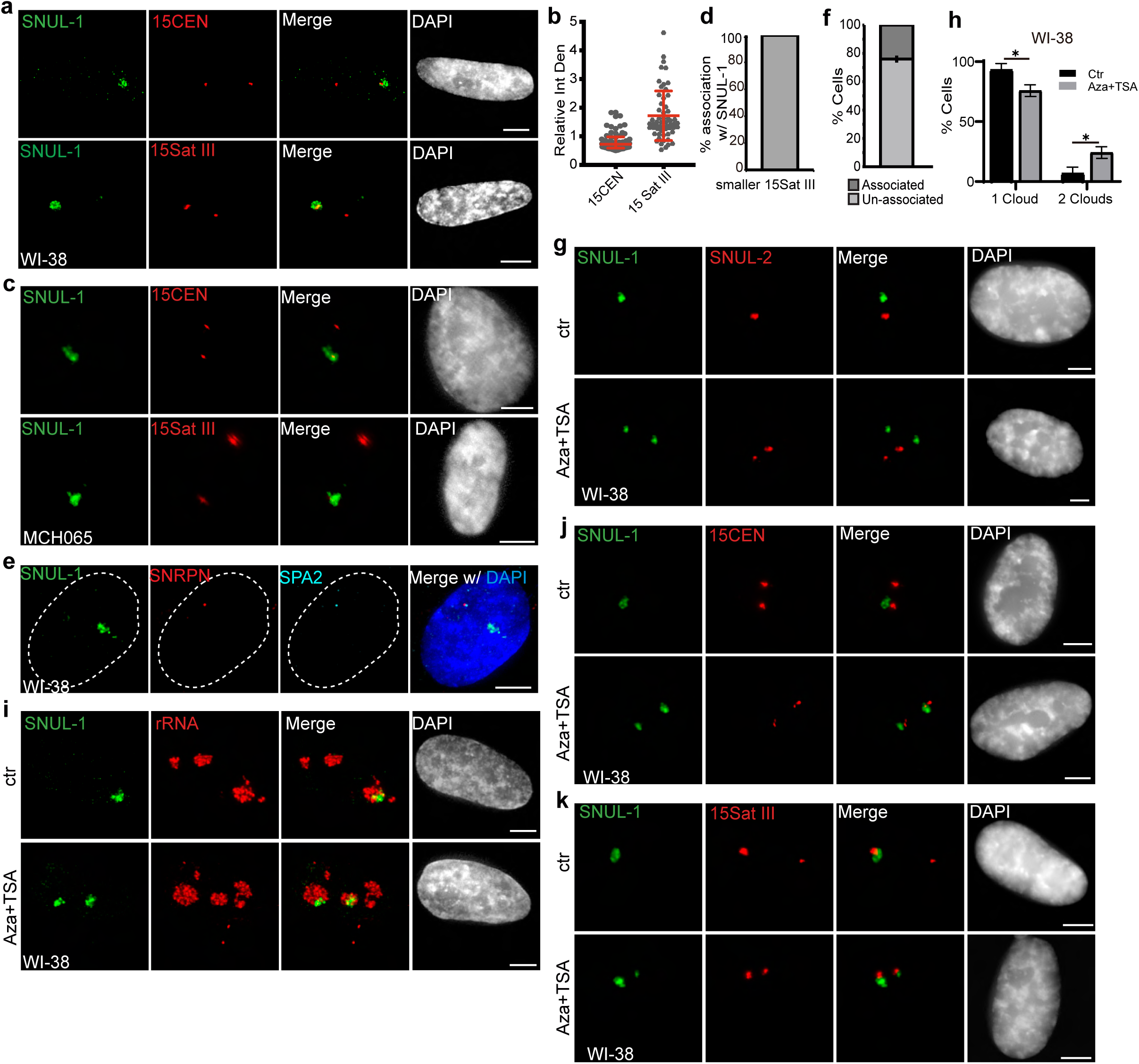
The SNUL-1 cloud displays mitotically-inherited random monoallelic association. **a**, Representative RNA-FISH images showing the association of SNUL-1 with the 15Sat III or 15CEN in WI-38 nuclei. **b**, Plot showing the relative integrated density of the 15Sat III signal in WI-38 nuclei. The relative integrated density is calculated by dividing the measurement of the larger DNA-FISH signal by that of the smaller DNA-FISH signal. Data are presented as Median and interquartile range. n = 60. **c**, Representative RNA-FISH images showing the association of SNUL-1 cloud with 15Sat III or 15CEN in MCH065 nuclei. **d**, Quantification of the association rates between SNUL-1 and the smaller 15Sat III in MCH065 nuclei. Data are presented as Mean ± SD from biological triplicates. > 50 cells were counted for each of the biological repeats. **e**, Representative RNA-FISH images showing the localization SNUL-1 along with the SNRPN and SPA2 transcription site on the paternal allele of Chr. 15 in WI-38 nucleus. Dotted lines mark the boundary of the nucleus. **f**. Quantification of the association rate between SNUL-1 and the transcription sites of SNRPN and SPA2 in WI-38 cells. Data are presented as Mean ± SD from biological triplicates. > 35 cells were counted for each of the biological repeats. **g**, Representative RNA-FISH images showing the distribution of SNUL-1 (green) and SNUL-2 (red) in control and Aza-dC (500nM) and TSA (80nM) treated WI-38 nuclei. **h**, Quantification of the percentage of cells showing one or two SNUL-1clouds in control and Aza+TSA-treated WI-38 cells. Data are presented as Mean ± SD from biological triplicates. > 50 cells were counted for each of the biological repeats. Student’s unpaired two-tailed t-tests were performed. *p < 0.05. **i**, RNA-FISH to detect SNUL-1 clouds in control and Aza+TSA-treated WI-38 nuclei. Nucleoli are visual- ized by rRNA (red). **j**, DNA-RNA-FISH of SNUL-1 RNA and 15CEN in control and Aza+TSA-treated WI-38 nuclei. **k**, DNA-RNA-FISH to detect SNUL-1 RNA and 15Sat III in control and Aza+TSA-treated in WI-38 nuclei. All scale bars, 5µm. DNA is counterstained by DAPI.

By utilizing the mouse A9 cells integrated with one allele of human Chr. 15 (mono- chromosomal somatic cell hybrid A9+H15)^18^, we further confirmed that SNUL-1 is indeed transcribed from Chr. 15 by RNA pol I and formed a confined RNA territory in the nucleolus. In the somatic-hybrid cells, the NOR on the transferred human chromosome remain silenced and showed no human rRNA expression, due to the inability of mouse-encoded RNA Pol I-specific transcription factors to bind to the human RNA pol I-transcribed gene promoters (Figure 3d; Ctr). Exogenous expression of human TBP-associated factors (TAF1A-D) in the A9+H15 cells reactivated RNA Pol I transcription from human Chr. 15, reflected by the presence of both human rRNA and SNUL-1 in the nucleolus (Figure 3d) ^18, 24^.

The rDNA content between the two alleles could vary profoundly in cell lines, as recently reported in the case of hTERT-RPE1 ^25^ (see also Figure S3g-h). Quantification of the integrated density of the rDNA spots on the mitotic chromosome spreads of WI-38 confirmed equal rDNA content between the two Chr.15 alleles (Figures 3e-f; green). This indicates that the monoallelic association of SNUL-1 to Chr. 15 is not dictated by the rDNA content in these cells. We further observed that the SNUL-1-associated Chr. 15 allele contained active NOR, as shown by positive 5-FU incorporation as well as the presence of RNA pol I transcription factor, UBF in the SNUL-1-decorated NORs (Figures S3i-j).

### SNUL-1 RNA displays mitotically inherited random monoallelic association (rMA) to the NOR of Chr. 15

We consistently observed a significant difference in the size of the Chr. 15-specific peri- centromeric Sat III repeat (15Sat III) signal between the two Chr. 15 alleles in multiple diploid cell lines ([WI-38; Figures 3e & 4a-b], [hTERT-RPE-1; Figures S3g & S4a-b], [MCH065; Figure 4c]), implying that these cells showed allele-specific differences in the amount or compaction of peri-centromeric 15Sat III DNA. Interestingly, in 100% of WI- 38 and hTERT-RPE1 cells, the SNUL-1 cloud was associated only with the larger 15Sat III signal containing Chr.15 allele (Figure 4a-b & S4a-b) (n =50 from biological triplicates). On the other hand, SNUL-1 cloud in the MCH065 cells was associated with the Chr. 15 allele containing the smaller 15Sat III signal (Figures 4c-d). These results imply that SNUL-1 non-randomly associate with a particular Chr. 15 allele in a cell type-specific manner. Loss-of-function studies revealed that SNULs did not influence allele-specific 15Sat III levels or compaction (Figure S4c-d).

We next determined whether SNUL-1 non-randomly associates with the paternal or maternal allele of the Chr. 15. Genes encoded within the imprinted Prader-Willi Syndrome (PWS)/Angelman Syndrome (AS) genomic loci ^26^, such as *SNRPN* and the lncRNA *SPA2*, are expressed only from the paternal allele of Chr. 15 ^27, 28^. In WI-38 cells (n=75), the SNUL-1 cloud was preferentially located away from the paternal Chr.15 allele, co-expressing *SNRPN* and *SPA2* (Figure 4e-f), indicating that in WI-38 cells SNUL-1 associated with the maternal allele of Chr. 15. On the other hand, in the MCH065 Fibroblasts and MCH065-derived iPSCs (MCH2-10), the SNUL-1 cloud was associated with the paternal Chr. 15 allele, as demonstrated by the localization of SNUL- 1 cloud next to the smaller 15SatIII or *SNRPN* RNA signals (Figures 4c-d & S4e-j). In all the tested cell lines (WI-38, hTERT-RPE1, and MCH2-10), the smaller 15SatIII signal was always associated with the paternal Chr. 15 (Figures S4i-j). Based on these results, we conclude that SNUL-1 is not an imprinted gene, but rather displays mitotically inherited random monoallelic association (rMA) to paternal or maternal Chr. 15 in a cell line-specific manner^29, 30^.

Repressive epigenetic modifiers control imprinted or monoallelic expression of lncRNAs ^31–33^. WI-38 cells incubated with DNA methyl transferase (DNMT; 5-Aza-2’- deoxycytidine [5-Aza-dC]) and histone deacetylase (HDAC; Trichostatin A [TSA]) inhibitors showed two separate SNUL-1 and SNUL-2 foci (Figure 4g-h). Both the SNUL-1 clouds in 5-Aza-dC+TSA-treated cells remained localized in the nucleolus (Figure 4i) and were associated with both Chr. 15 alleles in a population of cells (Figure 4j-k). These results indicate the potential involvement of repressive epigenetic regulators in maintaining the monoallelic association of SNULs.

### SNUL RNAs influence rRNA biogenesis

We next evaluated the potential involvement of SNUL territory in nucleolar functions. Iso-Seq and imaging data revealed that *SNUL-1* constituted a family of genes/transcripts sharing high sequence similarity. Modified antisense oligonucleotides (ASOs) targeting individual SNUL-1 CS candidate did not reduce total SNUL-1 levels (data not shown). The repeat sequence within SNULs was highly conserved among all the SNUL-1 CSs and was also shared by the SNUL-2 transcript. By using an ASO targeting this region (ASO-SNUL) we efficiently depleted both SNUL-1 and SNUL-2 (Figure 5a). Interestingly, SNUL-depleted cells showed enhanced 5-FU incorporation in the nucleolus (Figure 5b-c), and also showed increased levels of nascent 47S pre-rRNA, quantified by single molecule RNA-Fluorescent hybridization (smRNA-FISH) using a probe set (5’ETS-2) that preferentially detects nascent 47S pre-rRNA (Figures S5a-b). These results imply that SNUL depletion either enhanced the expression of nucleolus-enriched *rRNA* genes and/or reduced the co-transcriptional pre-rRNA processing. SNUL depletion did not alter the overall distribution of the nucleolus-localized proteins (Figures 5d & S5c-e) ^12, 34^. However, we observed that SNUL-depleted cells showed increased number of FC/DFC compartments/nucleolus, which could be a consequence of enhanced pre-rRNA levels in these cells (Figures 5d, 5f & S5c-d).

**Fig. 5.**
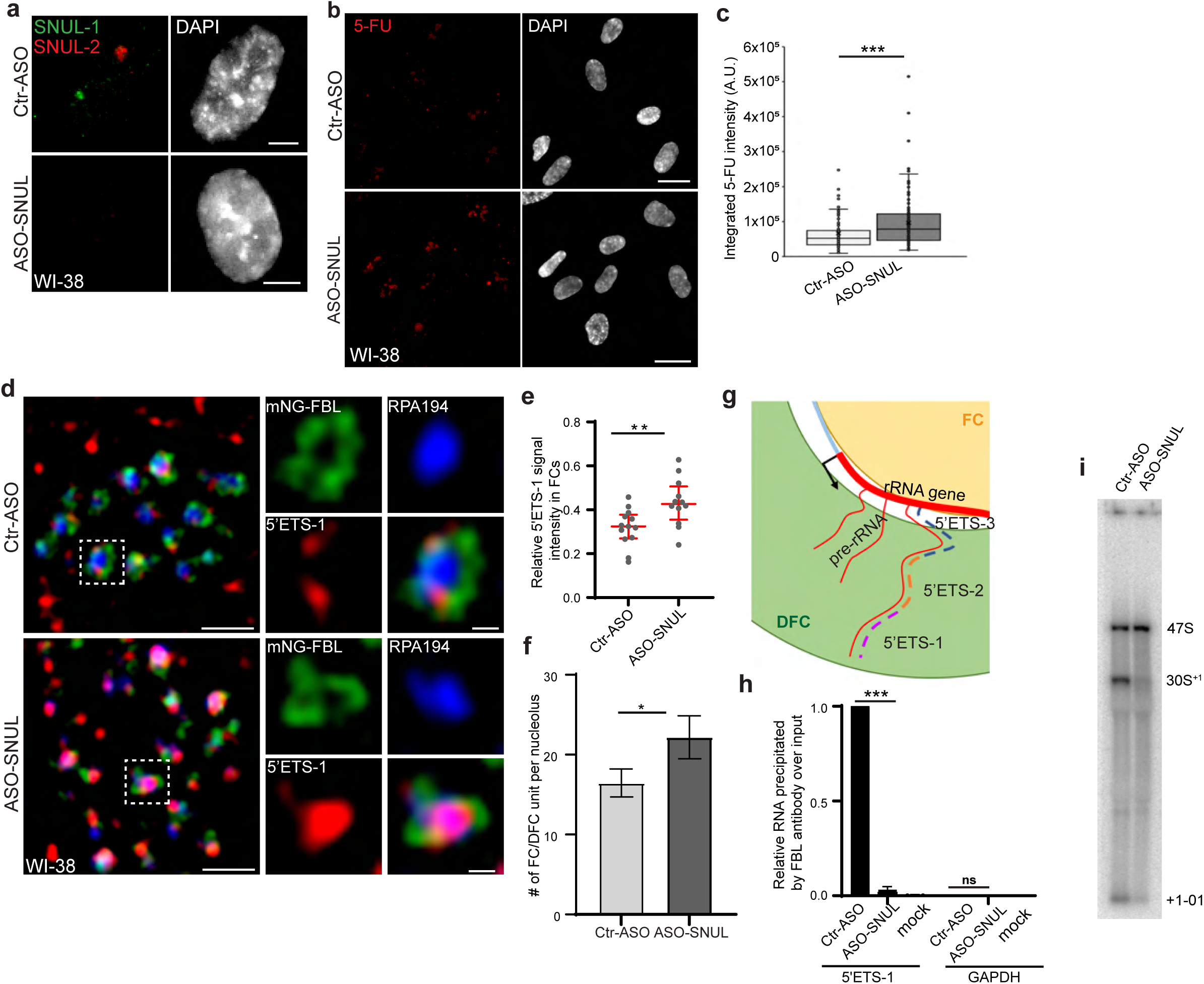
SNUL-1 influences rRNA biogenesis. **a**, RNA-FISH of SNUL-1 (green) and SNUL-2 (red) in WI-38 cells transfected with ctr-ASO or ASO-SNUL oligonucleotides. **b**, 5-FU immunostaining in control and SNUL-depleted WI-38 cells. Scale bars, 5 µm. Ctr-ASO and ASO-SNUL-treated cells were pulse labeled by 5-FU for 20 min. Scale bars, 20 µm. **c**, Boxplots of integrated 5-FU intensity per nucleus in control and SNUL-depleted WI-38 cells. Center line, median; box limits, upper and lower quartiles; whiskers, maximum or minimum of the data. Mann-Whitney test is performed. n =100. *p < 0.05, **p < 0.01, ***p < 0.001. **d**, SIM image of a single nucleolus showing the nascent pre-rRNA detected by 5’ETS-1 probe (red) in Ctr and SNUL-de- pleted cells. DFC is marked by mNG-FBL (green) and FC is marked by RPA194 (blue). Scale bars, 1 µm (main images) and 200nm (insets). **e**, Quantification of relative 5’ETS-1 intensity in FC of nucleolus in Ctr-ASO and ASO-SNUL cells. Center line, median. Mann-Whitney test is performed. **p < 0.01. >50 DFC/FC units from 10-15 nucleoli were counted for each sample. **f**, Quantification of the average number of FC/DFC unit per nucle- olus in control and SNUL-depleted WI-38 cells Data are presented as Mean ± SEM from nine biological repeats. > 15 nucleoli were counted from each experiment. *p < 0.05.**g**, Schematic showing the sorting of pre-rRNA in DFC/FC unit. The position of the 5’ETS regions are marked. **h**, Relative 5’ETS-1 precipitated by FBL antibody in control and SNUL-depleted WI-38 cells, ***p < 0.001. **i**, Northern blot using 5-ETS-1 probe from total RNA isolated from control and SNUL-depleted WI-38 cells.

The nascent 47S pre-rRNA is co-transcriptionally sorted from its transcription site at the DFC/FC boundary to DFC^21^. The DFC-localized FBL binds to the 5’ end upstream of the first cleavage site (01 site) (Figure S1m & 5g) of the 47S pre-rRNA co-transcriptionally and facilitates pre-rRNA sorting for efficient RNA processing and DFC assembly ^21^. Due to this, the 5’ end of 47S pre-rRNA is localized in the DFC region, as shown by SR-SIM of smRNA-FISH using the 5’ external transcribed spacer (5’ETS)-1 probe set targeting the first 414 nts of 47S pre-rRNA (Figure 5d) ^21^. On the other hand, the region within the 47S pre-rRNA 5’ETS located downstream of the 01 cleavage site (detected using 5’ETS-2 & 3 probes [Figures S1m & 5g; probes: 5’ETS-2 & 3]) associated with the rRNA transcription sites (FC or FC/DFC junction) (Figure S5c-d) ^21^. Interestingly, in the SNUL-depleted cells, FBL-interacting 5’ETS-1 region within the 47S pre-rRNA failed to sort to DFC, and instead preferentially accumulated in the FC (Figure 5d-e). Depletion of pre-rRNA processing factors, including FBL, compromised 47S pre-rRNA sorting at DFC, resulting in the accumulation of pre-rRNA in the FC region ^21^. Our results suggest the possibility that SNULs could influence pre-rRNA biogenesis by modulating FBL-mediated pre-rRNA sorting. In support of this, our SR-SIM imaging data revealed enriched association of SNUL-1 in the FBL-localized DFC. We therefore evaluated whether SNULs influence the interaction between FBL and pre-rRNA. Towards this, we performed FBL RNA-immunoprecipitation followed by quantitative RT-PCR to quantify the interaction between FIB and nascent pre-rRNA in control and SNUL-depleted cells. Strikingly, SNUL-depleted cells showed reduced association between FBL and pre-rRNA (Figure 5h), indicating that DFC-enriched SNULs could enhance the FBL interaction with pre-rRNA.

The sequence upstream of the 01-cleavage site within the 47S pre-rRNA (detected by 5’ETS-1 probe), is co-transcriptionally cleaved after it is sorted to DFC by FBL. Defects in the pre-rRNA sorting to DFC were shown to affect pre-rRNA processing ^21^. SNUL-1- depleted cells showed defects in the initial cleavage at the 5’end of the 47S pre-rRNA, as observed by the reduced levels of 30S+1 intermediate and +1-01 cleaved product (Figure 5i & S5f) by Northern blot analyses. The +1-01 is the unstable product processed from the 5’end of 47S pre-rRNA due to the cleavage at the 01 site. All these results indicate potential involvement of SNULs in pre-rRNA, sorting and/or co-transcriptional processing.

## DISCUSSION

We have discovered SNUL, a novel family of ncRNAs, which display non-random association to specific NOR-containing chromosomes within the nucleolus. Our data suggest that SNUL-1 is a member of a family of RNAs sharing similar sequence features. The most striking feature of the SNUL-1 sequence is its resemblance to the 21S pre- rRNA intermediate. Recent genomic mapping of acrocentric chromosome arms revealed that most of the sequences in the NOR-containing p-arms are shared among all the 5 chromosomes ^15, 18, 35^. However, these studies have also identified inter-chromosomal sequence variations ^18^. Our observations showing the association of individual members of SNULs to specific alleles of one of the acrocentric chromosomes support the idea that the acrocentric arms encode chromosome- and allele-specific transcripts.

The underlying mechanism(s) controlling differential expression rRNA gene copies in mammals is yet to be determined. Nucleolar dominance (NuD) is a developmentally regulated process that is speculated to act as a dosage-control system to adjust the number of actively transcribed rRNA genes according to the cellular need ^5, 36^. However, NuD is primarily observed in the ‘interspecies hybrids’ of plants, invertebrates, amphibians and mammals ^37–41^. NuD is reported in certain nonhybrid or ‘pure species’ of plants and fruit flies, but has not yet been observed in mammals, primarily due to lack of information about the allele-specific rRNA sequence variations ^42–44^. During NuD, chromosome allele-specific rRNA expression is observed, in which rRNA gene array within the NOR that is inherited from one parent (dominant) is maintained in a transcriptionally active status, while the rRNA loci from the other parent (under dominant) are preferentially silenced^38, 45–47^. Several features associated with the monoallelic regulation of SNULs share similarities with NuD ^37, 38^. SNUL-1, and also rRNA expression in NuD is dictated by non-imprinted random monoallelic expression. Repressive epigenetic modifiers play vital roles in the allele-specific expression of SNULs, and also rRNAs during NuD ^46, 48, 49^. Interestingly, specific sequence elements located near the NORs are implicated to control NuD ^50–52^. For example, in the germ cells of male *Drosophila*, Y chromosome sequence elements promote developmentally-regulated NuD of the Y chromosome- encoded rRNA over the X-chromosome rRNA^52^. It is possible that SNULs, by associating with an NOR allele could dictate allele-specific expression of rRNA arrays in human cells.

We observe that SNUL-1 forms a spatially constrained RNA territory that associates next to the NOR of Chr. 15, but is devoid of pre-rRNA. Furthermore, SNUL-depleted cells show elevated levels of pre-rRNA, along with defects in pre-rRNA sorting and processing. One-way SNUL-1 could modulate rRNA expression is via modulating the levels of bioprocessing machinery that control rRNA biogenesis and processing. For example, high sequence similarity between SNUL-1 and pre-rRNAs helps SNUL-1 to compete for and/or recruit factors that regulate rRNA biogenesis in a spatially constrained area within the nucleolus. A recent study, by visualizing the distribution of tagged pre-rRNAs from an NOR-containing chromosome, reported that similar to SNULs, pre-rRNAs transcribed from individual NORs form constrained territories that are tethered to the NOR-containing chromosomal regions ^53^. It is possible that SNUL-1, by forming a distinct RNA territory on the NOR of the Chr. 15 allele, influences the expression of rRNA genes from that NOR in an allele-specific manner. Such organization of SNULs and rRNA territories in a constrained area within the nucleolus would help to control the expression of a subset of rRNA genes without affecting the rRNA territories on other acrocentric chromosomes.

Our observation of compartmentalized distribution of individual members of SNUL RNA within specific sub-nucleolar regions challenges the current view that all the nucleoli within a single nucleus are composed of identical domains. Future work will entail determining the mechanism(s) underlying the constrained formation of ncRNA territories and allele-specific spreading and regulation on autosomal regions.

### Limitations of the present study

Presently, very little is known about the sequences in the short-arms of NOR-containing chromosomes, the region that harbors novel ncRNA genes such as SNULs. A recent study, by utilizing long-read sequencing in a haploid cell line revealed that p-arms of NOR-containing chromosomes are enriched with repeat sequences ^35^. Higher levels of sequence similarity observed between *SNUL-1* candidates and pre-rRNA made it impossible for us to precisely map the genomic coordinates of *SNUL-1* genes from the available long-read sequencing data set. Complete genome assembly of p-arms from SNUL-1-expressing diploid cells would be essential to map *SNUL-1* genes in the genome and also to identify the regulatory elements controlling monoallelic expression of *SNULs*. Genomic annotation of the full-length *SNUL-1* genes is also crucial for designing strategies to specifically alter the expression of individual *SNUL* genes, without targeting other SNUL-like genes, furthering mechanistic understanding of SNUL functions. Even with these technical limitations, the current study is highly impactful because our observations of the association of autosome arms by SNULs supports a paradigm-shifting model that ncRNA-coating of chromosomes and their roles in gene repression are not restricted only to sex chromosomes. In addition, our study will serve as a starting point towards the understanding of how differential rDNA expression is achieved during physiological processes. Altogether, this study will form the basis for an entirely new avenue of investigations, which would help to understand the role of ncRNAs on monoallelic changes in autosomal chromatin structure and gene expression in the nucleolus.

## ACKNOWLEDGEMENTS

We thank members of Prasanth’s laboratory and Dr. Ashish Lal (NCI, NIH) for their valuable comments. We thank Drs. Eric Bolton (UIUC) for sharing prostate cancer cell lines (PC3 and LNCap), Dr. Alok Sharma and Dr. Drinda Swanson from Abbott Inc. for CEN15 and 15Sat III DNA probes, Prof. Ling-ling Chen (SIBCB) for lentiviral constructs expressing fluorescently-labeled nucleolar proteins and Prof. Kyosuke Nagata (University of Tsukuba) for SL1 plasmids. We thank Dr. Jason Underwood (PacBiosciences) for the help to perform Iso-Seq. This work was supported by National Institute of Health R21-AG065748 & R01-GM132458 to KVP, GM099669, GM125196 to SGP, R35GM131819 to SS, R21AG065748 and R01GM123314 to SCJ, Cancer center at Illinois seed grants and Prairie Dragon Paddlers to KVP, National Science Foundation (NSF) to KVP [EAGER, 1723008], SGP [career award, 1243372 & 1818286] and SCJ [1940422 &1908992]. H.J. acknowledges support from the NIH (R01-GM120552). SMF is an employee of Ionis Inc., and ET and JK work for PACBIO and receive salary from the respective companies.

## MATERIALS AND METHODS

### Cell Culture

WI-38 and IMR-90 cells were grown in MEM medium supplemented with 10% fetal bovine serum (FBS), non-essential amino acid, sodium-pyruvate. HeLa and U2OS cells were grown in DMEM medium supplemented with 5% FBS. hTERT-RPE1, mouse A9+H15 and SH-SY5Y cells were grown in DMEM/F12 medium supplemented with 10% FBS. GM12878 cells were grown in RPMI1640 medium supplemented with 15% FBS. PC-3 cells were grown in RPMI-1640 medium supplemented with 10% FBS. MDA-MB-231 and LNCaP cells were grown in RPMI 1640 media supplemented with 10% FBS. SaOS-2 cells were grown in McCoy’s 5A medium supplemented with 15% FBS. MCF10A cells were grown in DMEM/F12 medium supplemented with 5% house serum, hydrocortisone, cholera toxin, insulin, and EGF. HS578 cells were grown in DMEM medium supplemented with 10% FBS and insulin. MCH065 cells were grown in DMEM medium supplemented with 10% FBS. All media were supplemented with Penicillin/Streptomycin. H9 hESCs and MCH2-10 iPSCs were grown on acid-treated coverslips coated with Matrigel® hESC-Qualified Matrix (Corning®, Product Number 354277) in mTeSR™ Plus (STEMCELL Technologies™, Catalog #100-0276). Cells were maintained in a 5% CO_2_ incubator at 37 ℃. Cell lines are obtained from commercial vendors such as ATCC and Coriell. Cell lines used in our study has been authenticated by STR profiling (UIUC Cancer Center). All cell lines were checked for mycoplasma.

### Transfection and virus infection

For ASO treatments, Ctr-ASO or ASO-SNUL were transfected to cells at a final concentration of 100 nM using Lipofectamine RNAiMax Reagent (Invitrogen). Cells were cultured for another 3 days before harvest.

pHAGE-mNG-C1-FBL and pHAGE-mTagBFP2-C1-B23 plasmids were gifts from Dr. Ling-ling Chen’s lab ^21^. HeLa cells in 3.5 cm dish were transfected with 500ng of pHAGE-mNG-C1-FBL and/or pHAGE-mTagBFP2-C1-B23 using Lipofectamine 3000 Reagent (Invitrogen).

pCHA-hTAF1A-D plasmid were gifts from Dr. Kyosuke Nagata’s lab ^24^. A9+H15 cells in 3.5 cm dish were transfected with pCHA-hTAF1A-D plasmid, 400 ng per each plasmid, using Lipofectamine 3000 Reagent (Invitrogen).

For stably expressing mNG-FBL in WI-38 cells, lentiviral particles were packaged by transfecting pHAGE-mNG-C1-FBL, pMD2.G, psPAX2 to 293T cells in WI-38 growing medium. Virus were collected twice at 48 h and 72 h after transfection, removed of cell debris by centrifuge, and snap frozen. WI-38 cells were infected by the virus for 2 days and changed back to medium without virus.

### Transcription Inhibition and Epigenetic Marker Inhibition

For RNA Pol I inhibition, cells were treated with 1 μM BMH21 (Selleckchem), 10 ng/ml ActD (Sigma-Aldrich), or 1μM CX5461 (Sigma-Aldrich) for 45 min or 2 h. For RNA Pol II inhibition, cells were treated with 1) 5 μg/ml ActD for 2 h, 2) 2.5 μM Flavopiridol (Selleckchem) for 3 h, or 3) 32 μg/ml DRB for 3 h. After 3 h of DRB treatment, cells were washed with PBS for 3 times and were recovered in fresh growth medium for 30 min or 60 min. For epigenetic mark inhibition, cells were treated with 80nM TSA and 500nM 5-Aza-dC for 6 days.

### RNA-Fluorescence *in situ* hybridization (FISH)

For all of the FISH and Immunofluorescence staining done with adherent cells, cells were seeded on #1.5 coverslips at least two days before experiments. For GM12878 and isolated HeLa nucleoli, suspension was smeared onto the Poly-L-lysine-coated (Sigma-Aldrich) coverslips prior to fixation.

For RNA-FISH using probes prepared by nick translation, cells were fixed by 4% PFA for 15 min at room temperature (rt) and permeabilized with 0.5% Triton X-100 for 5 min on ice. Alternatively, cells were pre-extracted by 0.5% Triton X-100 in CSK buffer for 5 min on ice and then fixed by 4% PFA for 10 min. Probes were made using Nick Translation Kit (Abbott Molecular) as per manufacturer’s instructions, added to the hybridization buffer (50% formamide, 10% dextran sulfate in 2XSSC supplemented with yeast tRNA), and before hybridization. Hybridization was carried out in a humidified chamber in the dark overnight at 37 ℃. The coverslips were then washed in 2X SSC and 1X SSC and 4X SSC. DNA is counterstained with DAPI. Coverslips were mounted in VectaShield Antifade Mounting Medium (Vector Laboratories) or ProLong Diamond Antifade Mountant (Invitrogen). Please see Table S4 for primer and probe sequence details.

5’ETS smFISH probe sets were described in ^21^. SNRPN and SPA2 smFISH probe sets were designed using Stellaris^®^ Probe Designer. Oligonucleotides with 3’ amino group (LGC Biosearch Technologies) were pooled and coupled with either Cy^®^3 Mono NHS Ester (GE healthcare) or Alexa Fluor^TM^ (AF) 647 NHS Ester (Invitrogen) by incubation overnight at 37 ℃ in 0.1M NaHCO_3_. Probes were then purified by G-50 column (GE Healthcare) and ethanol precipitation. Concentration was measured by the OD at 550nm (Cy^®^3) or 650nm (AF647). For RNA-FISH involving smFISH probes, smFISH probes were added to Stellaris^®^ RNA FISH Hybridization Buffer (LGC Biosearch Technologies) with 10% formamide at a final concentration of 125nM. Hybridization was carried out in a humidified chamber in the dark for 6 h at 37 ℃. The coverslips were then washed with Stellaris^®^ RNA FISH Wash Buffer A and mounted as described above.

Digoxin-labeled RNA probes were in vitro transcribed as per manufacturers’ instructions (DIG RNA labeling Mix, Roche; T7 polymerase, Promega; SP6 Polymerase, Promega) and purified by G-50 column (GE Healthcare). For RNA-FISH using ribo- probes, cells on coverslips were fixed by 4% PFA for 10 min at rt, and then treated with 0.25% acetic anhydride in 0.1 M triethanolamide (pH 8.0) for 10 min. Coverslips were washed in 1XSSC for 5min, treated with 0.2N HCl for 10 min, and pre-hybridized in 50% formamide, 5XSSC for at least 6 h at rt. Dig-labeled RNA probes were added to the hybridization buffer (50% formamide, 5XSSC, 1X Denhardt’s solution, 0.1% Tween20, 0.1% [w/v] CHAPS, 100 μg/ml Heparin, 5 mM EDTA, and 50 μg/ml Yeast tRNA) at a final concentration of 2 μg/ml. Hybridization was carried out in a humidified chamber in the dark overnight at 50 ℃. The coverslips were then washed with 0.2XSSC for 1 h at 55 ℃, blocked in 4% BSA, PBS for 30 min at 37 ℃, and incubated with anti-Dig-FITC or - Rhodamine (1:200) (Roche) in 1% BSA, PBS for 1 h at 37 ℃. The coverslips were washed twice with washing buffer (0.1% Tween20, 2XSSC) and refixed with 4% PFA for 15 min at rt.

For RNase A treatment, pre-extracted cells were incubated with 1 mg/ml RNase A in CSK buffer for 30 min at 37 ℃. Cells were then fixed by 4% PFA for 15 min at rt and processed to RNA-FISH. For DNase I treatment, fixed and permeabilized cells were incubated with 200 U/ml DNase I (Sigma) in DNase I buffer prepared with PBS for 2 h at 37 ℃, followed by incubation in Stop solution for 10 min at room temperature. RNA-FISH was then performed as described above.

### RNA-DNA FISH

For DNA-FISH using chromosome paint probes (Chrs. 13, 15, 22) (MetaSystems), after fixation and permeabilization, coverslips were incubated in 20% glycerol overnight and then went through freeze-thaw by liquid nitrogen for at least 6 cycles. Coverslips were then treated with 0.1N HCl for 5 min and prehybridized in 50% formamide, 2XSSC for 30 min at rt. Probe mix was made by adding the RNA-FISH probe into the chromosome paint probe. Probes were applied to the coverslips and denatured with the coverslips at 75-80 ℃ on a heating block. Hybridization was carried out in a humidified chamber in the dark for 48 h at 37 ℃.

For FISH using DNA-FISH probes made by nick translation, cells were pre-extracted and fixed. Salmon sperm DNA and Human Cot-I DNA were added to the hybridization buffer. Denaturation and hybridization were performed as described above.

### DNA-FISH on Metaphase Spread

Cells were grown to ∼70% confluence and treated with KaryoMax Colcemid solution (Gibco) at a final concentration of 0.1 μg/ml in growth medium for 3 h. Mitotic cells were then shaken off and pelleted by centrifuge. Cells were then gently resuspended in 75 mM KCl and incubated at 37 ℃ for 30-40 min. Cells were then fixed by freshly prepared fixative (methanol: acetic acid 3:1 [v/v]) and dropped onto pre-cleaned microscope slides from height. After air-drying, slides were stored at -20 ℃ for a least overnight before the DNA-FISH. For the DNA-FISH on metaphase chromosomes, slides were rehydrated with PBS and then treated with 50 μg/ml Pepsin in 0.01N HCl at 37 ℃ for 9 min. Slide were then rinsed by PBS and 0.85% NaCl sequentially and dehydrated by a series of Ethanol at different concentration (70%, 90%, and 100%). Air-dried slides were then subjected to hybridization as described above.

### Immunofluorescence staining (IF)

Cells on coverslips were fixed and permeabilized before blocking in 1% BSA for 30 min at rt. Coverslips were then incubated with primary antibodies (anti-FBL, 1:500, Novus Biologicals, NB300-269; anti-RPA194, 1:50, Santa Cruz, sc-48385; anti-UBF, 1:50, Santa Cruz, sc-13125; anti-DDX21, 1:20000, proteintech,10528-1-AP). and secondary antibody (anti-mouse IgG2a AF647, 1:2000, Invitrogen, A-21241; anti-mouse IgG AF568, 1:2000, Invitrogen, A-11031; anti-mouse Cy5, 1:1000), sequentially. Coverslips were then washed with PBS and refixed with 4% PFA. RNA-FISH was then carried out if needed.

### 5-FU metabolic labeling

Cells were grown to ∼70% confluence on the day of experiments. Cells were treated with 2 mM 5-FU (Sigma-Aldrich, F5130) for specified time periods before harvest. To detect incorporated 5-FU, IF was performed with anti-BrdU antibody (1:800, Sigma-Aldrich, B9434) as described above.

### Nucleoli isolation

HeLa nucleoli were isolated as described in ^54^ with adjustments. Briefly, HeLa cells were collected by trypsinization and lysed in nuclear extraction buffer (50 mM Tris-HCL, pH7.4, 0.14 M NaCl, 1.5 mM MgCl_2_, 0.5% NP-40, 1mM DTT, and RNase Inhibitor). Nuclei was precipitated and resuspended in S1 solution (0.25 M sucrose and 10mM MgCl_2_). Nuclear suspension was gently layered on S2 solution (0.35 M sucrose and 0.5 mM MgCl_2_) and spun at 2,000g for 5 min at 4 ℃. Purified nuclei were then sonicated by Bioruptor UCD-200 at high mode until nucleoli were released. Another sucrose cushion was then carried out with S3 solution (0.88 M sucrose and 0.5 mM MgCl_2_). Isolated nucleoli were then resuspended in S2 and subjected to RNA extraction by Trizol Reagent (Invitrogen) or RNA-FISH.

### Northern Blot

For the pre-rRNA Northern, 2 μg of total RNA extracted from WI-38 cells treated with Ctr-ASO or ASO-SNUL was separated on 1% denature agarose gel prepared with NorthernMax Denaturing Gel Buffer (Ambion) and run in NorthernMax Running Buffer (Ambion). RNA was then transferred to Amersham Hybond-N+ blot (GE Healthcare) by capillary transfer in 10 x SSC and crosslinked to the blot by UV (254 nm, 120mJ/cm2). The DNA probes were labeled with [α-32P] dCTP by Prime-It II Random Primer Labeling Kit (Stratagene) as per manufacturer’s instructions. Hybridization was carried out using ULTRAhyb Hybridization Buffer (Ambion) containing 1 X 10^6^ cpm/ml of denatured radiolabeled probes overnight at 42 ℃. Blots were then washed and developed using phosphor-imager.

### Native RNA Immunoprecipitation

WI38 cells were washed twice with cold PBS and collected by centrifuge (1,000g, 10 mins at 4 ℃). Cells were then lysed in 2ml RIP buffer (50 mM Tris pH 7.4, 150 mM NaCl, 0.05% Igepal, 1 mM phenylmethyl sulfonyl fluoride (PMSF), 1μM Leupeptin, 1μM Pepstatin, 0.2 μM Aprotinin, and 2mM VRC(NEB)), and sonicated by Bioruptor UCD-200 at high mode on ice. Cells were centrifuged at 1,000g at 4 ℃ for 10 mins and supernatants were then pre-cleared with 15 μl Dynabeads Protein G (Invitrogen) for 30 mins. FBL antibody (Abcam) or rabbit IgG2b was incubated with 25μl Dynabeads Protein G for 30 mins. The cell supernatants were then incubated with Dynabeads Protein G at 4 ℃ for 2hrs. The Dynabeads Protein G were then washed with high salt buffer (50 mM Tris pH 7.4, 650 mM NaCl, 0.15% Igepal, 0.5% sodium deoxycholate,1 mM phenylmethyl sulfonyl fluoride (PMSF), 1μM Leupeptin, 1μM Pepstatin, 0.2 μM Aprotinin, and 2mM VRC(NEB)) for three times and RIP buffer twice, followed by RNA isolation and RT-qPCR.

### Imaging Acquisition

For widefield microscopy, z-stack images were taken using either 1) DeltaVision microscope (GE Healthcare) equipped with 60X/1.42 NA oil immersion objective (Olympus) and CoolSNAP-HQ2 camera, or 2) Axioimager.Z1 microscope (Zeiss) equipped with 63X/1.4 NA oil immersion objective and Zeiss Axiocam 506 mono camera. Images were then processed through deconvolution and maximum intensity projection.

SIM images were taken using either DeltaVision OMX V3 system (GE Healthcare) equipped with a 100X/1.4 NA oil immersion objective, 3 laser beams (405nm, 488nm, and 568nm) and EMCCD camera (Cascade II 512), or SR-SIM Elyra system (Zeiss) equipped with 63X/1.4 NA oil immersion objective, 4 laser beams (405nm, 488nm, 561nm, and 642nm). For DeltaVision OMX V3 system, Channels were aligned for each of the experiments using the registration slide and TetraSpeck microspheres (Invitrogen).

SIM image stacks were acquired with a z-interval of 0.125 μm, 5 phases and 3 angles. SIM reconstruction and registration of channels were performed by softWoRX software (GE Healthcare). For SR-SIM Elyra system, channels were aligned for each of the experiments using the TetraSpeck microspheres (Invitrogen). SIM image stacks were acquired with a z-interval of 0.125 μm, 5 phases and 3 rotations. SIM reconstruction and channel alignment were performed by ZEN 2011 software (Zeiss).

### Imaging Analyses

For colocalization analyses, 3D SIM stacks were imported into Fiji/ImageJ. The nucleolar area containing SNUL-1 signal was selected and Pearson’s correlation coefficients (no threshold) were calculated by the Coloc2 Plugin.

For the measurement of integrated density, z-stacks were imported into Fiji/ImageJ and maximum intensity projection was performed. Signal of interest was then segmented by Maximum Entropy Multi- Threshold function in the ij-Plugins Toolkit with number of thresholds = 3. A Binary mask was generated based on the second level of the threshold from the last step. The integrated density of the signal of interest from the original image within the mask was then measured. For rDNA contents on the two Chr. 15 alleles in WI38 cells, relative integrated density was calculated by dividing the measurement of the rDNA signal on the Chr. 15 with the larger 15Sat III by the measurement on the other Chr. 15. For rDNA contents on the two Chr. 15 alleles in hTERT-RPE1 cells, relative integrated density was calculated by dividing the measurement of the larger rDNA signal by the measurement of the smaller rDNA signal. For 15Sat III and 15CEN, relative integrated density was calculated by dividing the measurement of the larger signal spot by that of the other spot signal within the same cell.

For the measurement of 5-FU incorporation and 5’ETS-2 signal in control and SNUL- depleted cells, z-stacks were imported into Fiji/ImageJ and maximum intensity projection was performed. Nuclei were segmented by optimized threshold and inverted into binary mask. The integrated density of 5-FU signal or 5’ETS-2 signal in each of the nuclei was measured.

For the measurement of 5’ETS-1 signal intensity in FC of nucleolus in control and SNUL-depleted cells, the middle z session was selected for each image and imported into Fiji/ImageJ then split into single channels. FCs were segmented by auto threshold of RPA194 channel and inverted into binary mask. The binary masks were applied to the 5’ETS-1 channel and the integrated intensity of 5’ETS-1 signals within FCs were counted for each nucleolus. The relative 5’ETS-1 signal intensity in FCs was calculated by dividing the integrated intensity of 5’ETS-1 signals within FCs by the integrated intensity of the entire image.

### PacBio Iso-Seq

Total RNA from isolated HeLa nucleoli was poly-adenylated by Poly(A) Polymerase Tailing Kit (Epicentre) and depleted of rRNA by the RiboMinus™ Human/Mouse Transcriptome Isolation Kit (Invitrogen). RNA was then reverse transcribed by the SMARTer PCR cDNA Synthesis Kit (Clontech) and amplified for 15 cycles using KAPA HiFi PCR Kit (KAPA biosystems). cDNA was then separated into two fractions by size using 0.5X and 1X AMPure PB Beads (Pacific Scientific), respectively. SNUL-1 was then enriched from the two fractions by xGen capture procedure with SNUL-1 Probe 4 using the xGen hybridization and Wash Kit (IDT). Another round of PCR amplification was carried out after the capture. The two fractions were then combined. Library was prepared by Amplicon SMRTbell Prep (Pacific Scientific) and sequenced on LR SMRT cell with 20 h movie.

### Nanopore sequencing

Total RNA from isolated HeLa nucleoli was depleted of rRNA by the RiboMinus™ Human/Mouse Transcriptome Isolation Kit (Invitrogen). RNA was converted to double stranded cDNA using random hexamer with the NEBNext Ultra RNA First Strand and NEBNext Ultra RNA 2^nd^ Strand Synthesis Kits (NEB). 1D library was prepared with the SQK-LSK108 kit (Oxford Nanopore) and sequenced on a SpotON Flowcell MK I (R9.4) flowcell for 14 h using a MinION MK 1B sequencer. The flowcell was washed and another identical library was loaded in the same flowcell and sequenced for another 14 h. Basecalling was performed with Albacore 2.0.2.

The PacBio Iso-seq and nanopore RNA-seq data sets are deposited to the NCBI SR data base. The Bioproject accession number is SRA data: PRJNA814414.

### Sequencing Analyses

To find transcripts that are similar to SNUL in the high-quality PacBio database, RIBlast^55^, a tool for predicting RNA-RNA interactions, is used in its default setting for SNUL-1 probe 1 as the query RNA. The top 2000 candidates with the lowest interaction energy were intersected to generate the final set of 507 transcripts similar to SNUL-1. Pairwise BLASTs of each of the top ranked PacBio Iso-Seq clones and genome assembly hg38 were performed to pick the candidates showing the least similarity with any of the annotated genes.

A reference fasta file was generated with the 5 picked candidates from PacBio sequencing dataset. The fast5 file from nanopore long read sequencing was basecalled using Guppy. The obtained fastq file containing 1267135 reads (across two runs) was aligned to the hg38 reference fasta file using minimap2 ^56^. Alignment statistics were computed using samtools – flagstat option ^57^, a total of 123582 (9.7%) were mapped to the reference fasta file. Mapped reads were extracted into a sam file and indexed using samtools, for visualizing the alignments through IGV (Integrative Genomics Viewer) ^58^. To generate an accurate version of SNUL-1 transcripts, we generated a consensus sequence from the long-read alignments, using samtools and bcftools ^59^. To evaluate the specificity of the assembled transcripts, we performed a similarity comparison between the generated consensus sequence against rRNA and PacBio Iso-Seq CS clones. We observed that the sequences identified/generated from our analysis were more analogous to the Iso-Seq clones over rRNA.

In order to verify the error rate of PacBio sequencing technology, for each isoform in the high-quality PacBio database, we ran BLAST against the human transcript database (GRCh38.p13 assembly). The maximum number of target sequences and the maximum number of high-scoring segment pairs were set to 20 and 1 respectively, and the rest of the arguments were set to default in the BLAST runs. LAGAN-v2.0 ^60^ was then used to perform pairwise global alignment between each isoform and its corresponding top 20 best matches, found by BLAST. A dissimilarity score was assigned to each matched candidate by taking the ratio of mismatching sites to all the sites where both isoform and the matched candidate did not contain gaps. The matched candidate with the least dissimilarity score was taken as the best match to the isoform. The mean of the dissimilarity scores, associated with isoforms having GC content greater than 60% (matching the GC content of ITS1), being ∼0.5% verifies the PacBio error rate (<1%).

To determine whether the 5 isoforms capable of detecting SNUL-1 are different transcripts, and their difference is not due to sequencing error, we propose the following three hypotheses to be tested:

*H_0_*: There is one known gene whose transcripts are *I_1_,…,I_5_*.

*H_1_*: There is one unannotated gene, i.e., with no transcripts present in human transcript database, whose transcripts are *I_1_,…,I_5_*.

*H_2_*: There are multiple unannotated/annotated genes whose transcripts are *I_1_,…,I_5_*.

If *H_0_* is true, then there exists a known transcript such that the dissimilarity score between isoform *i* (*I_i_*) and the transcript’s dissimilarity score should follow the empirical distribution of the dissimilarity scores in Figure S1l. For each isoform, the empirical probability of observing a dissimilarity score greater than or equal to its associated dissimilarity score was computed (Table S2). The product of the empirical p-values being in the order of 10^-10^ suggests that *H_0_* does not hold.

If *H_1_* is true, there should be one unannotated transcript whose dissimilarity score with each read is about 0.5% (the empirical mean of the dissimilarity scores). Therefore, the pairwise dissimilarity scores for the isoforms should be about 1%. We computed the real pairwise dissimilarity by doing global pairwise alignment for each pair of isoforms using LAGAN (Table S3) . The pairwise dissimilarities being 4% or above suggest that *H_1_*is not true. Approximating the empirical distribution of the dissimilarity scores with an exponential probability density function with mean 0.5%, if *H_1_*holds then the pairwise dissimilarity should follow erlang distribution with shape and scale parameters being 2 and 200 respectively, as the sum of two independent exponential random variables with the same rate parameter has erlang distribution. The probability of observing a pairwise dissimilarity score greater than or equal to each real pairwise dissimilarity score under erlang distribution was computed. The product of these probabilities being in the order of 10^-39^ rejects the *H_1_* hypothesis which leaves us with accepting *H_2_* hypothesis.

For the analyses of alignment between SNUL-1 candidates and pre-rRNA, the candidate sequences were aligned to the canonical 21S sequence and to each other using the LAST algorithm as described previously ^61^.

### Data analyses and statistics

The data used in this study are performed at least biological triplicates. Statistical analyses (two-tailed Student’s t-test and Mann-Whitney test) were done by GraphPad Prism. Please see figure legends for details about the sample sizes and statistical significance.

## SUPPLEMENTARY TABLES

**Table S1. Sequence of the SNUL-1 probe** TATGGCTCCT TCCTCCCTCT CTCCATTCTT CTCTCAGCTT TCCTGTGGGC 50 AGGGGTAGGC ACAGCCAGGC TTGGGAGCAT CGCCATGCCC TGCCACCTGG 100 GTCCCAGCCT GCTCCTCGTT ATAGTCTTCC CAGTTTGGGG AAGAGCAGTG 150 ATATGCCAAG AATGGAGGCC TCAGACTCTC CCAATCCCTG ATTTTTACAT 200 GTCCCCCTAT AAGGCCCCTC TGCCATCTAC ACTTTTGCCC TTCATCCACA 250 AAGCCCAAAA GGAAGGCATT ATAGCTAGCC ATGCCCTCTG ACTGCCCTCT 300 GCCCCTTTAA GGGAAATGGA AATGGGTACC CAGCTGACTG AACCTACTCA 350 ACACCTCCAG AAATTAGACA CTAGGGCATG GTGCCACCCT CCCAGGCTGG 400 CACATGCTAC CCTGGCAGAG GATCAAATAA CCCCCCCATC ATACCCTGCC 450 CCATGTCTTC CTCTACTCTC TCCCTCATGC TTTCTCTCTC TCTCTCTCTC 500 TCTGTCTCTC TCTCTCTCTC TCTCTCAGCT CAAAGCACAG CTGAGCCTTA 550 AAAGGGGGGT TGAGGGGGTG GAGAGACCAA GCTGGGGCAG GGGGGTATAG 600 AGCTCCAATA GCACGTTTTC ACCT

**Table S2. The CS candidate sequences.**

Please see the Excel file

**Table S3.** The dissimilarity score between each isoform and its best match from human transcript database.

| isoform name | dissimilarity score (%) | empirical p-value |
| --- | --- | --- |
| nucDNA1_PK_combo__HQ_transcript/1269 | 5.5 | 0.02 |
| nucDNA1_PK_combo__HQ_transcript/2305 | 4.2 | 0.02 |
| nucDNA1_PK_combo__HQ_transcript/2615 | 1.2 | 0.05 |
| nucDNA1_PK_combo__HQ_transcript/9572 | 3.3 | 0.03 |
| nucDNA1_PK_combo__HQ_transcript/8644 | 4.3 | 0.02 |

For each isoform the empirical p-value is the empirical probability of observing a dissimilarity score greater than or equal to its dissimilarity score. The product of the 5 p-values being in the order of $10^{-10}$ rejects the hypothesis of all isoforms being the transcripts of the same known gene.

**Table S4.** The pairwise dissimilarity for the isoforms.

| **Isoform 1** | **Isoform 2** | **pairwise dissimilarity (%)** | **p-value** |
| --- | --- | --- | --- |
| transcript/2615 | transcript/1269 | 4.8 | 0.0005 |
| transcript/2615 | transcript/2305 | 4.8 | 0.0004 |
| transcript/2615 | transcript/9572 | 6.4 | 2E-05 |
| transcript/2615 | transcript/8644 | 6.6 | 2E-05 |
| transcript/1269 | transcript/2305 | 4.0 | 0.002 |
| transcript/1269 | transcript/9572 | 5.2 | 0.0002 |
| transcript/1269 | transcript/8644 | 4.2 | 0.001 |
| transcript/2305 | transcript/9572 | 5.0 | 0.0004 |
| transcript/2305 | transcript/8644 | 5.1 | 0.0003 |
| transcript/9572 | transcript/8644 | 4.3 | 0.001 |

Global pairwise alignment was used to compute the dissimilarity score for each pair of isoforms. The dissimilarity score was computed by taking the ratio of mismatching to matching sites where both isoforms do not contain gaps. For each pair of isoforms, the p-value shows the probability of observing a dissimilarity score greater than or equal to their dissimilarity by approximating the empirical distribution of pairwise dissimilarity with erlang distribution.

**Table S5. Probes used in this study.**

Please see the Excel file.

---

**Extended Data Fig. 1.**
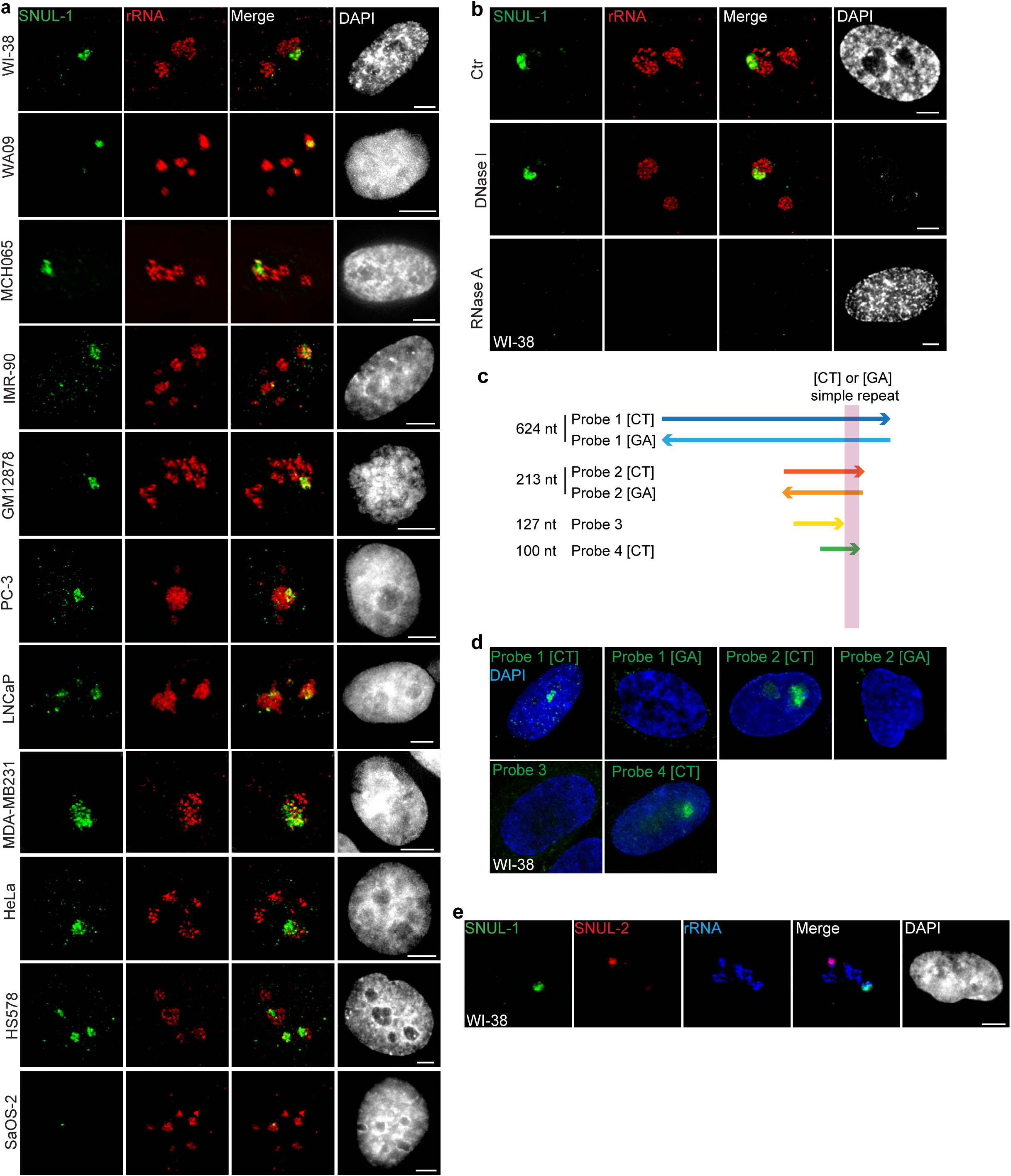
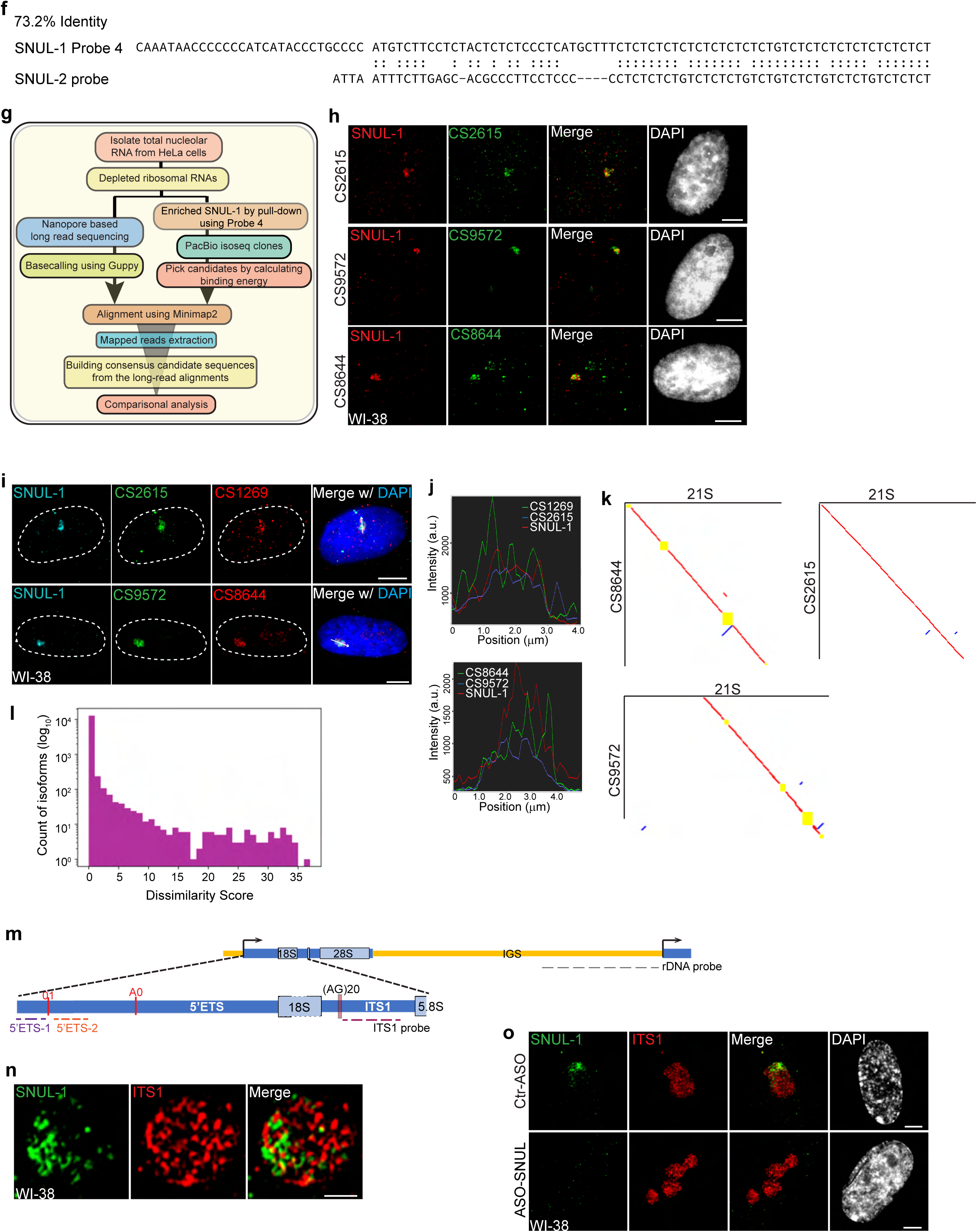
SNUL-1 forms RNA clouds in human cell lines. **a**, RNA-FISH of SNUL-1 (green) in different human cell lines. For all images, nucleoli are visualized by rRNA (red). Scale bars, 5µm. **b**, RNA-FISH of SNUL-1 after nuclease treatments in WI-38 cells. Scale bars, 5µm. **c**, Schematic showing the truncated probes designed to determine the minimum region required for SNUL-1 hybridization. **d**, RNA-FISH performed with the strand-specific ribo-probes listed in c. Scale bars, 5µm. **e**, RNA-FISH to detect SNUL-1 (green) and SNUL-2 (red) clouds in WI-38 cells. Nucleoli are visualized by rRNA (blue). Scale bars, 5µm. **f**, Local alignment between SNUL-1 Probe 4 and SNUL-2 probe. Note the imperfect [CT] repeat in SNUL-2 probe and the poor alignment between the two probes beyond the [CT]-rich region. **g**, Schematic showing the workflow of the unbiased strategies to determine the full-length SNUL-1 sequence. **h**, RNA-FISH using probes designed from the CSs (green) and SNUL-1 (red) Probes 1 in WI-38 cells. Scale bars, 5µm. **i**, RNA-FISH with CS probes and SNUL-1 Probe 1 in WI-38 cells. Scale bars, 5µm. **j**. Signal profiles of the lines marked in i. Note that the signals shown by different probes are not completely colocalized. **k**, Pairwise sequence comparisons between 21S and CS8644, 21S and CS2615, and 21S and CS9572. Red lines indicate forward aligned regions, blue lines indicate reverse aligned regions, and yellow boxes indicate unaligned regions. **l**, Histogram of the dissimilarity score between each isoform in the high-quality PacBio database and its best match in the human transcript database. **m**, Schematic showing the positions of the rRNA and ITS1 probes. **n**, Representative SIM image showing the relative distribution of SNUL-1 (green) and pre-rRNA hybridized by ITS1 probe (red) within a single nucleolus. Scale bars, 1µm. **o**, RNA-FISH of SNUL-1 (green) and pre-rRNA hybridized by ITS1 probe (red) in WI-38 cells transfected by Ctr-ASO or ASO-SNUL. Scale bars, 5µm. DNA is counterstained with DAPI.

**Extended Data Fig. 2.**
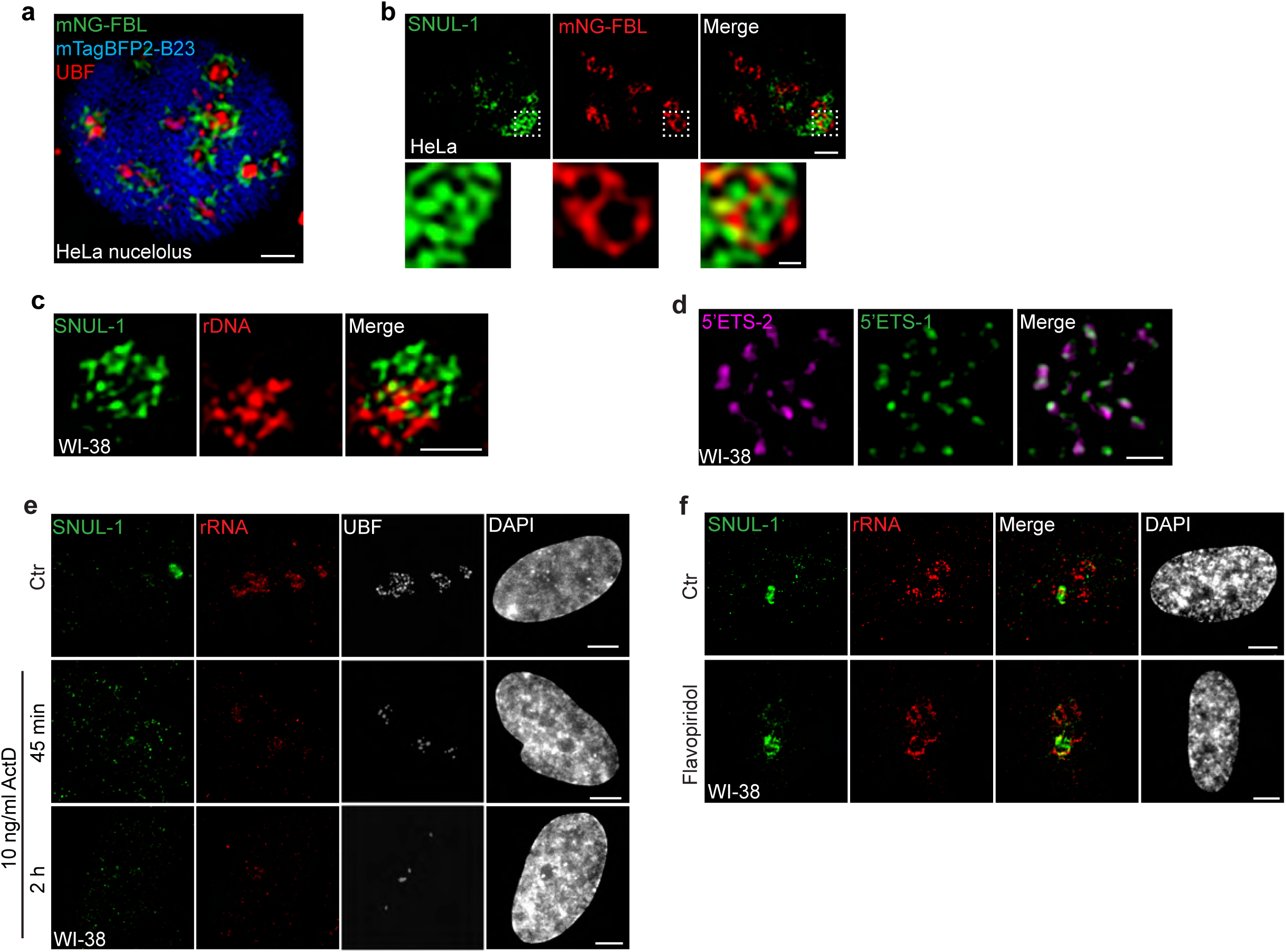
SNUL-1 is an RNA Pol I transcript and forms constrained nucleolar territory. **a**, Visualization of the tripartite structure within a single HeLa nucleolus by SIM. FC is marked by UBF (red), DFC is marked by mNeonGreen (mNG)-FBL (green), and GC is marked by mTagBFP2-B23 (blue). Scale bar, 1 µm. **b**, Representative SIM image of a single nucleolus showing the SNUL-1 distribution relative to DFC/FC units in HeLa cells. DFC is marked by mNG-FBL. Scale bars, 1µm (main images) and 200nm (insets). **c**, Representative SIM image showing the relative distribution of SNUL-1 (green) and rDNA (red) within a single nucleolus. Scale bars, 1 µm. Note: The prominent signal of rDNA represents clusters of inactive rDNA repeats. **d**, Representative SIM image of a single nucleolus showing the distribution of nascent pre-rRNA detected by 5’ETS-1 and 5’ETS-2 probes. Scale bars, 1 µm. **e**, Co-RNA-FISH and IF to detect SNUL-1 (green), rRNA (red) and UBF (white) in control and RNA pol I-inhibited (low con. Of ActD) WI-38 cells. Scale bars, 5 µm. **f**, RNA-FISH to detect SNUL-1 (green), rRNA (red) in control and RNA pol II-inhibited WI-38 cells. Scale bars, 5µm. DNA is counter- stained with DAPI.

**Extended Data Fig. 3.**
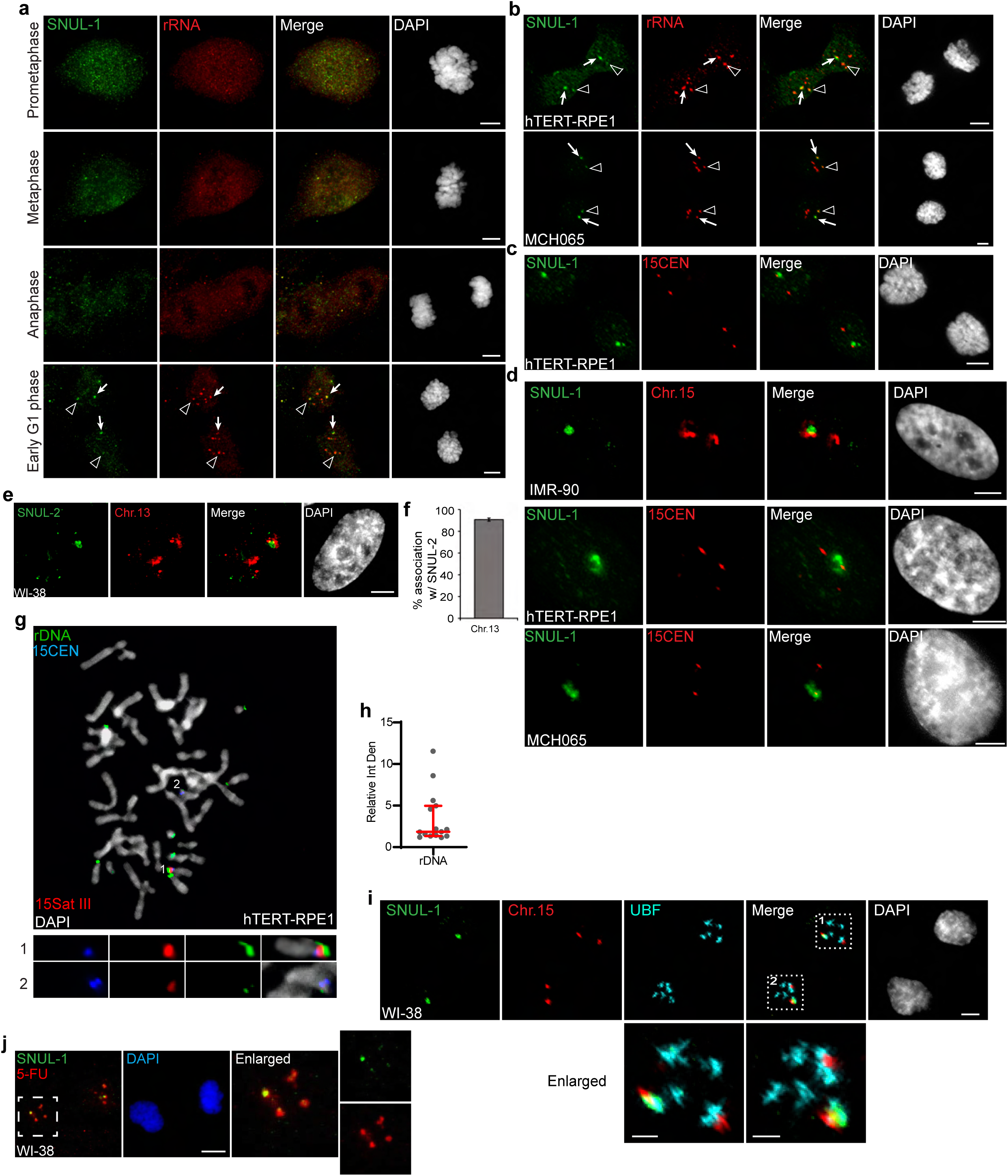
SNUL-1 is associated with the NOR of one Chr. 15 allele. **a**, RNA-FISH showing the distribution of SNUL-1 in WI-38 cells during mitosis. Arrows point at the prominent SNUL-1 cloud in early G1 daughter nuclei. Arrow heads point at the relatively weak SNUL-1 cloud in early G1 phase of daughter nuclei. Scale bars, 5 µm. **b**, RNA-FISH of SNUL-1 and rRNA in early G1 daughter nuclei. Arrows point at the prominent SNUL-1 clouds. Arrow heads point at the relatively weak SNUL-1 clouds. Scale bars, 5 µm. **c**, DNA-RNA-FISH to detect SNUL-1 RNA and 15CEN in early G1 hTRET-RPE1 daughter nuclei. Scale bars, 5 µm. **d**, DNA-RNA-FISH to detect SNUL-1 RNA and Chr.15 in the nucleus. The two alleles of Chr.15 are marked by either probe painting the q-arms of the chromosome (IMR-90 cells), or 15CEN (hTRET-RPE1 and MCH065 cells). **e**, DNA-RNA-FISH to detect SNUL-2 RNA and Chr. 13 marked by the probe painting the q-arm of the chromosome in WI-38 nucleus. Scale bars, 5 µm. **f**, Quantification of the association rates between SNUL-2 and Chr13. Data are presented as Mean ± SD from biological triplicates. > 100 cells were counted for each of the biological repeats. **g**, DNA-FISH showing the rDNA contents on Chr.15 in hTRET-RPE1 metaphase chromosomes. The two alleles of Chr.15 are marked by 15Sat III and 15CEN. rDNA arrays are detected by a probe within the IGS region. **h**, Relative integrated density of the rDNA contents on the two Chr.15 rDNA arrays is calculated by dividing the measurement of the larger rDNA signal by the smaller rDNA cloud. Data are presented as Median and interquartile range. n = 15. **i**, Immuno-RNA & DNA-FISH showing SNUL-1 (green), Chr. 15 alleles (red) and UBF (blue) in early G1 phase WI-38 daughter nuclei. Scale bars, 5 µm (main images) and 2 µm (insets). **j**, SNUL-1 localization and 5-FU incorporation in WI-38 telophase/early G1 daughter nuclei. Scale bars, 5 µm. DNA is counterstained with DAPI.

**Extended Data Fig. 4.**
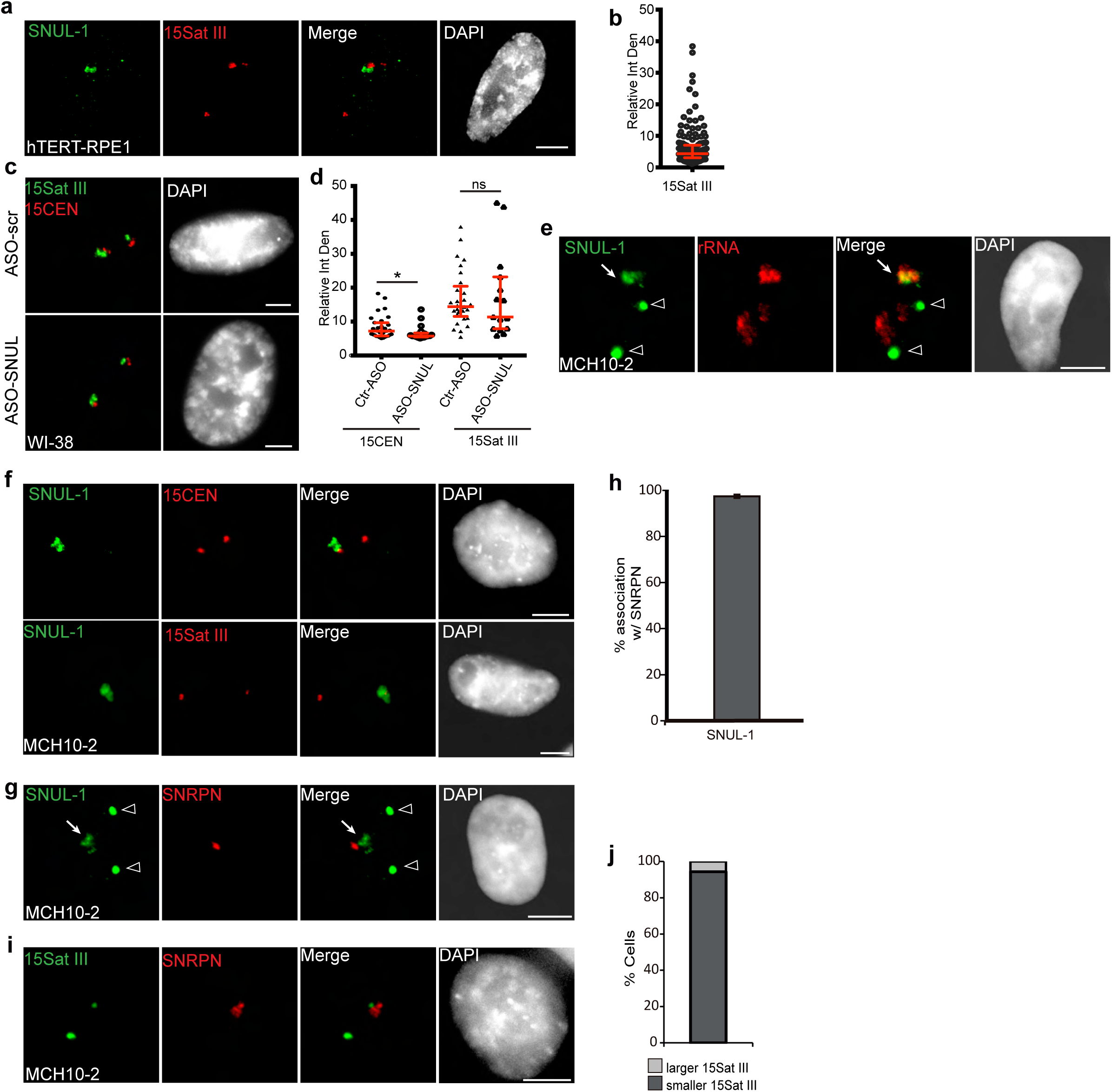
The SNUL-1 cloud displays mitotically-inherited random monoallelic association. **a**, DNA-RNA-FISH showing the localization SNUL-1 RNA cloud and 15Sat III in hTERT-RPE1 cell nucleus. **b**, Plot showing the relative integrated density of the 15Sat III signals. Relative integrated density is calculated by dividing the score of the larger 15Sat III signal by that of the smaller 15Sat III signal. Data are presented as Median and interquartile range. n = 149. **c**, DNA-FISH to detect 15Sat III and 15CEN in Ctr and SNUL-depleted WI-38 nuclei. **d**, Plot showing the relative integrated density of the 15Sat III or 5CEN signals in control and SNUL-depleted WI-38 cells. Relative integrated density is calculated by dividing the measurement of the larger DNA-FISH signal by that of the smaller DNA-FISH signal. Data are presented as Median and interquartile range. n = 30. Mann-Whitney tests are performed. *p < 0.05; ns, not significant. **e**, RNA-FISH showing the distribution of SNUL-1 and rRNA in MCH2-10 nuclei. Arrows point at the SNUL-1 cloud. Arrowheads mark two non-nucleolar foci of unknown origin hybridized by the SNUL-1 probe only in MCH2-10 nuclei. **f**, DNA-RNA-FISH of SNUL-1 RNA and 15CEN or 15Sat III in MCH2-10 IPSC nuclei. Please note that the SNUL-1 probe-hybridized non-nucleolar foci is observed only after RNA-FISH and not after RNA-DNA-FISH treatments. **g**, Representative RNA-FISH image showing the localization of SNUL-1 cloud and SNRPN transcription site in MCH2-10 IPSC nucleus. Arrows point at the SNUL-1 cloud. Arrowheads mark two non-nucleolar foci of unknown origin hybridized by the SNUL-1 probe only in MCH2-10 nuclei. **h**, Quantification of the association rates between SNUL-1 and SNRPN in MCH2-10 IPSCs. Data are presented as Mean ± SEM from biological triplicates. > 50 cells were counted for each of the biological repeats. **i**, Representative DNA-RNA-FISH image to detect SNRPN RNA and 15Sat III in MCH2-10 nucleus. **j**, Quantification of the association rates between SNRPN RNA signal and the smaller and larger 15Sat III in MCH2-10 cells. > 100 cells were counted. All scale bars, 5µm. DNA is counterstained with DAPI.

**Extended Data Fig. 5.**
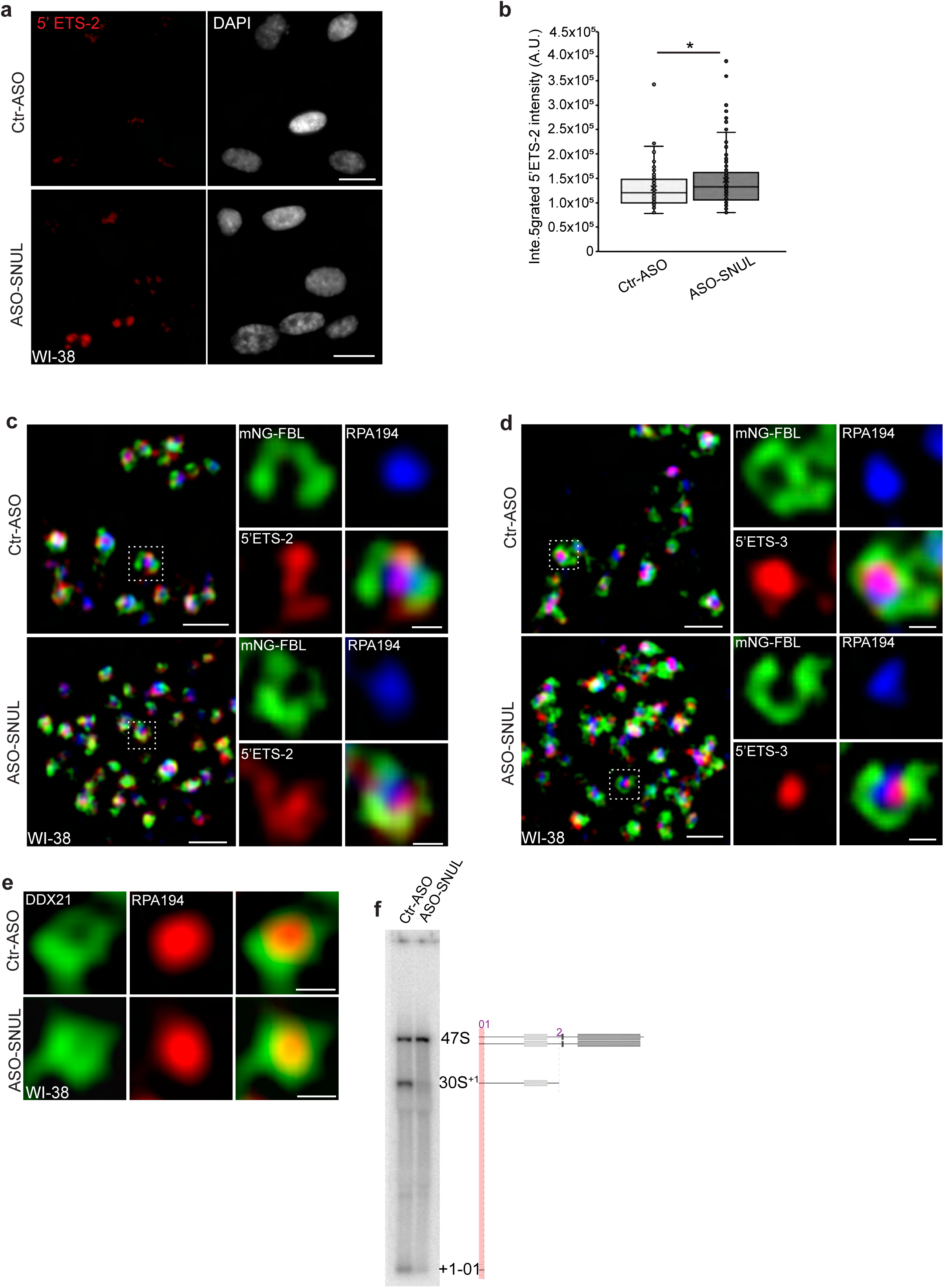
SNUL-1 influences rRNA biogenesis. **a**, Representative RNA-FISH images showing nascent pre-rRNA levels (detected by 5’ ETS-2 probe) in control and SNUL-depleted cells. Scale bars, 20 µm. **b**, Boxplots of integrated 5’ETS-2 intensity per nucleus in Ctr-ASO and ASO-SNUL treated cells. Center line, median; box limits, upper and lower quartiles; whiskers, maximum or minimum of the data. Mann-Whitney test is performed. n = 100 *p < 0.05. **c**, SIM image of a single nucleolus showing the nascent pre-rRNA detected by 5’ETS-2 probe (red) in Ctr and SNUL-depleted cells. DFC is marked by mNG-FBL (green) and FC is marked by RPA194 (blue). Scale bars, 1 µm (main images) and 200nm (insets). **d**, SIM image of a single nucleolus showing the nascent pre-rRNA detected by 5’ETS-3 probe (red) in Ctr and SNUL-depleted cells. DFC is marked by mNG-FBL (green) and FC is marked by RPA194 (blue). Scale bars, 1 µm (main images) and 200nm (insets). **e**, SIM images of a single FC/DFC unit showing the localization RPA194 and DDX21 in Ctr and SNUL-depleted cells. Scale bars, 200 nm. **f**, Northern blot (presented in figure 5i showing the schematics of the pre-rRNA species that are detected by 5’ETS-1 probe in control and SNUL-depleted cells.

